# Interruption of the Intratumor CD8:Treg Crosstalk Improves the Efficacy of PD-1 Immunotherapy

**DOI:** 10.1101/2023.05.15.540889

**Authors:** Shannon N Geels, Alexander Moshensky, Rachel S Sousa, Benjamin L Walker, Rima Singh, Giselle Gutierrez, Michael Hwang, Thorsten R Mempel, Qing Nie, Shivashankar Othy, Francesco Marangoni

**Affiliations:** Institute for Immunology, University of California Irvine, Irvine CA; Department of Physiology and Biophysics, University of California Irvine, Irvine CA; Center for Complex Biological Systems, University of California Irvine, Irvine CA; NSF-Simons Center for Multiscale Cell Fate Research, University of California Irvine, Irvine CA; Department of Molecular Biology and Biochemistry, University of California Irvine, Irvine CA; Center for Immunology and Inflammatory Diseases (CIID), Massachusetts General Hospital, Boston MA; Harvard Medical School, Boston MA; Department of Developmental and Cell Biology, University of California Irvine, Irvine CA

**Author notes:** Correspondence should be addressed to F.M. These contributors share senior authorship (Shivashankar Othy, Francesco Marangoni).

**Keywords:** T regulatory (Treg) cells, Programmed cell death protein 1 (PD-1), Tumor tolerance, Functional Intravital Microscopy (F-IVM)

## Abstract

PD-1 blockade unleashes the potent antitumor activity of CD8 cells but can also promote immunosuppressive T regulatory (Treg) cells, which may worsen response to immunotherapy. Tumor Treg inhibition is a promising strategy to overcome therapeutic resistance; however, the mechanisms supporting tumor Tregs during PD-1 immunotherapy are largely unexplored. Here, we report that PD-1 blockade increases tumor Tregs in mouse models of immunogenic tumors, including melanoma, and metastatic melanoma patients. Unexpectedly, Treg accumulation was not caused by Treg-intrinsic inhibition of PD-1 signaling but instead depended on an indirect effect of activated CD8 cells. CD8 cells colocalized with Tregs within tumors and produced IL-2, especially after PD-1 immunotherapy. IL-2 upregulated the anti-apoptotic protein ICOS on tumor Tregs, causing their accumulation. ICOS signaling inhibition before PD-1 immunotherapy resulted in increased control of immunogenic melanoma. Thus, interrupting the intratumor CD8:Treg crosstalk is a novel strategy that may enhance the efficacy of immunotherapy in patients.

## INTRODUCTION

Cancer immunotherapy using checkpoint blockade has revolutionized the management of previously intractable malignancies by extending overall and progression-free survival in people with a wide range of metastatic cancers (André et al., 2020; Borghaei et al., 2015; Ferris et al., 2016; Larkin et al., 2019; Migden et al., 2018; Motzer et al., 2018; Nghiem et al., 2016). Checkpoint blockade removes the mechanisms that limit the activation of effector T cells, unleashing a strong response against tumors. PD-1 inhibition is the foundation of most checkpoint immunotherapy strategies; however, the majority of patients either do not respond to this treatment or relapse (Sharma et al., 2017). Discovering the mechanisms underlying treatment failure is a crucial prerequisite for designing new, more efficacious antitumor strategies based on PD-1 antagonism.

Engagement of PD-1 on activated T cells by PD-L1 and PD-L2 recruits Shp2 and other phosphatases to the immunological synapse, in turn suppressing TCR and CD28 signaling (Hui et al., 2017; Yokosuka et al., 2012). The rationale behind PD-1 immunotherapy is to unleash the antitumor function of effector T cells, especially CD8 cells (Sade-Feldman et al., 2018; Tumeh et al., 2014; Wei et al., 2017), by antibody-mediated interruption of PD-1 interaction with its ligands. However, the impact of PD-1 blockade on tumor immunity may extend far beyond the stimulation of effector T cells. αPD-1 antibodies likely alter the information flow among several immune cell types in the tumor environment via direct PD-1 binding, by instructing the production of soluble mediators of immunity, or both. It is therefore paramount to understand the effects of PD-1 blockade on the tumor immune environment as a whole, including its potential to trigger immunosuppressive mechanisms that limit therapeutic efficacy.

Among the various immunosuppressive cells within the tumor microenvironment, we focused our investigation on CD4^+^ Foxp3^+^ T regulatory (Treg) cells. Tregs respond to tumor-associated antigens in secondary lymphoid organs by upregulating chemokine receptors such as CCR4, CXCR3, and CCR5 necessary for recruitment to non-lymphoid tissues, including tumors (Marangoni et al., 2018). Tregs reencounter their cognate antigen during brief interactions with dendritic cells (DCs) in the tumor microenvironment (Marangoni et al., 2021) and consequently instruct local immune suppression (Bauer et al., 2014). Accordingly, Treg accumulation in tumors is an adverse prognostic factor in multiple cancers, including melanoma (Shang et al., 2015).

While widespread Treg depletion would facilitate tumor rejection, it also poses the risk of developing severe autoimmunity. Thus, it is critical to understand how PD-1 immunotherapy modulates tumor Treg responses, so we may locally disable their immunosuppressive function. Tumor Tregs express PD-1, and PD-1 blockade may support Treg numbers and activation in gastro-esophageal cancer (Kamada et al., 2019). Further, higher PD-1 expression in Treg compared to CD8 cells predicts failure of checkpoint immunotherapy (Kumagai et al., 2020). However, the mechanisms by which PD-1 inhibition supports tumor Tregs remain to be studied.

Here, we investigated the causes of Treg expansion after PD-1 blockade using patient data and mouse models, Treg-specific deletion of critical genes, intravital microscopy, and intercellular communication analysis via CellChat (Jin et al., 2021). We found that cancer immunogenicity is a crucial factor in increasing tumor Treg numbers during PD-1 blockade and that such expansion limits the efficacy of immunotherapy. Unexpectedly, Treg accumulation was not due to cell-intrinsic inhibition of PD-1 signaling in Tregs; instead, αPD1-mediated activation of CD8 cells promoted Treg increase. The intratumor CD8:Treg crosstalk was mediated by IL-2 and ICOS. Administration of αICOSL antibodies acted as an immune conditioning regimen for the tumor environment, which increased the effectiveness of subsequent PD-1 immunotherapy.

## RESULTS

### Tumor immunogenicity drives Treg accumulation during PD-1 blockade

We started to investigate the mechanisms supporting Tregs after PD-1 blockade by considering tumor immunogenicity. PD-1 blockade may increase the activation of self-reactive Tregs in non-immunogenic tumors. Immunogenic tumors may also support Tregs since they can respond to an inflamed environment or neoantigens (Ahmadzadeh et al., 2019). Therefore, we compared T cell populations in mouse models of non-immunogenic and immunogenic melanoma treated with αPD-1 or isotype control antibodies. We chose melanoma because it is sensitive to PD-1 blockade, yet more than half of the patients do not respond to immunotherapy (Larkin et al., 2019). We used the D4M.3A cell line isolated from BrafV600E / PTEN-null mice (Jenkins et al., 2014) since these strong driver mutations give rise to a poorly immunogenic tumor (Lo et al., 2021). On the other hand, a D4M derivative expressing the SIINFEKL peptide from chicken ovalbumin (D4M-S) is highly immunogenic and sensitive to PD-1 immunotherapy (Di Pilato et al., 2019). To focus our investigation on immune responses within the tumor environment, we treated established melanomas with PD-1 immunotherapy in the presence of FTY720, an S1PR1 functional antagonist that blocks lymphocyte egress from lymph nodes (Fig. 1A). We found that Treg, CD8, and CD4^+^Foxp3^−^ T helper (Th) cells poorly accumulated in non-immunogenic D4M melanomas; numbers and percentages of these cells were not altered by αPD-1 treatment. Conversely, immunogenic D4M-S tumors had more CD8, Th, and Treg cells than D4M melanomas at baseline, and PD-1 blockade significantly enhanced CD8 and Treg abundance (Fig. 1B-D, S1A-C). Administration of αPD-1 increased the proliferation of CD8 cells as estimated by Ki67 expression (Fig. 1E, F, and S1D), as well as IFNγ and TNF production (Fig. S1E). Granzyme B expression remained unchanged (Fig. S1F). These data are consistent with other reports claiming that CD8 cells are the primary target of PD-1 immunotherapy (Sade-Feldman et al., 2018; Wei et al., 2017). Notably, PD-1 inhibition did not significantly increase Ki67 expression in Tregs (Fig. 1E, F, and S1D) but resulted in the upregulation of Foxp3 and the activation markers GITR and ICOS (Fig. 1G). Similar results were obtained using the immunogenic MC38 colon carcinoma (Fig. S1G-I). Altogether these data show that PD-1 inhibition leads to elevated Treg numbers, not accompanied by increased proliferation, and to increased expression of activation markers in two distinct immunogenic tumor models.

**Figure 1.**
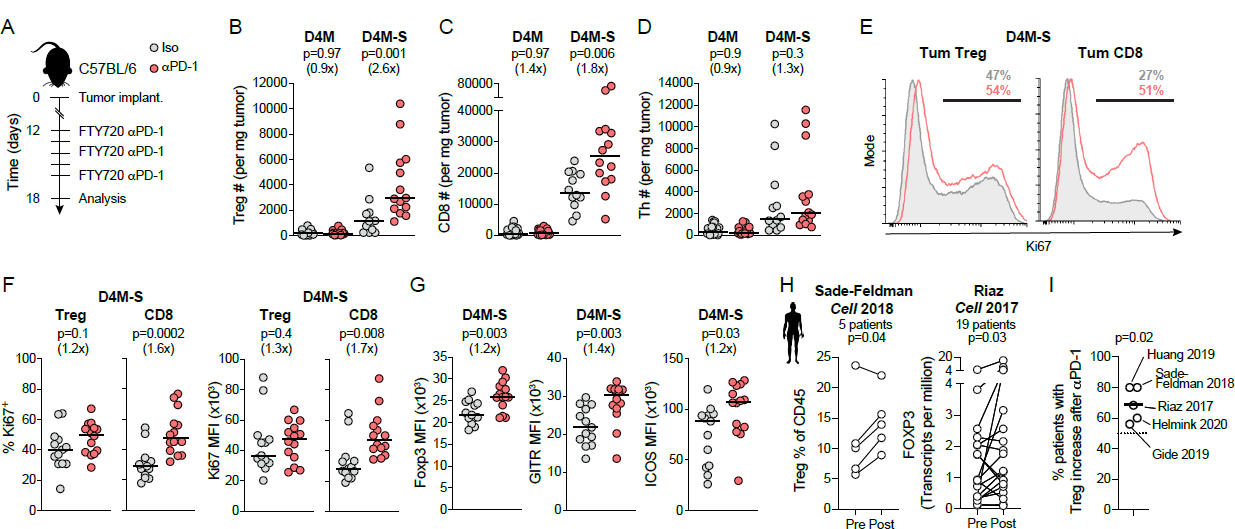
PD-1 blockade locally increases tumor Treg counts. A. Experimental scheme to assess the impact of PD-1 blockade on tumor-associated T cells. **B-D.** Tumor Treg (**B**), CD8 (**C**), and Th (**D**) cell numbers per mg of tumor in mice bearing D4M or D4M-S melanomas treated or not with αPD-1. **E, F.** Representative histograms (**E**) and quantification of Ki67 expression (**F**) in Treg and CD8 cells in D4M-S tumors treated or not with αPD-1. **G.** MFI of Foxp3, GITR, and ICOS in tumor Tregs with or without αPD-1. For **B-G**, n=25 (D4M) and 12-14 (D4M-S) mice/group pooled from 5 (D4M) or 3 (D4M-S) independent experiments. Bars depict the median values of the distribution. p values by Mann-Whitney U test. **H.** Treg quantification in single-cell (Sade-Feldman) and bulk (Riaz) RNA sequencing datasets. p values by paired Student’s *t*-test. **I.** Comparison of patients with increased Tregs after PD-1 blockade in the available single-cell RNA sequencing (Sade-Feldman), bulk RNA sequencing (Riaz, Helmink, Gide), and immunofluorescence (Huang) datasets. p value by one-sample *t*-test against the theoretical value of 50%, corresponding to no increase. The solid bar depicts the mean value.

We then studied how PD-1 blockade activates Tregs in tumor-draining lymph nodes. In secondary lymphoid organs, Tregs exist in a resting state characterized by the CD44^lo^CD62L^+^ phenotype (“central” or cTregs), and an activated state identified by the CD44^hi^CD62L^−^ phenotype (“effector” or eTregs) (Smigiel et al., 2014). We found that the numbers of lymph node Treg, CD8, and Th cells increased with αPD-1 treatment, irrespective of tumor immunogenicity. The percentage of CD8 and Th cells was unchanged. Still, there was a trend towards increased proportions of Tregs after PD-1 blockade (Fig. S1J-L), possibly due to the PD-1-mediated restriction of lymph node Treg activation at homeostasis (Pereira et al., 2023). We also observed a tendency for eTregs and activated CD44^hi^CD62L^−^ CD8 cells to accumulate after PD-1 blockade (**Fig. S1M**), accompanied by increased proliferation (Fig. S1N). PD-1 blockade did not change the expression of Foxp3, GITR, and ICOS in lymph node eTregs (Fig. S1O). Thus, unlike in tumors, PD-1 inhibition induces lymph node Treg proliferation without increased expression of activation markers. Moreover, we found that eTregs were heavily skewed towards PD1-deficient cells in radiation chimeras reconstituted with a 1:1 mixture of WT and PD-1^-/-^ bone marrow. Notably, the percentage of PD-1^-/-^ cells did not further increase in the tumor (Fig. S1P, Q), corroborating the notion that the consequences of PD-1 inhibition are different in lymph node compared to tumor Treg cells.

### PD-1 immunotherapy increases Tregs in human melanoma

To extend our observations to humans, we conducted a meta-analysis of tumor Tregs in metastatic melanoma patients treated with PD-1 monotherapy. We compiled publicly available datasets encompassing single-cell RNA sequencing (Sade-Feldman et al., 2018), bulk RNA sequencing (Gide et al., 2019; Helmink et al., 2020; Riaz et al., 2017), and immunofluorescence (Huang et al., 2019). We reanalyzed these data to compare Treg levels in the same patients before and after the administration of PD-1 monotherapy. Paired analysis of Tregs revealed a statistical increase in the single-cell RNA sequencing dataset (Sade-Feldman et al., 2018) and in one of the bulk RNA sequencing datasets (Riaz et al., 2017) (Fig. 1H). Treg increase did not reach statistical significance in the other two bulk RNA sequencing datasets (Gide et al., 2019; Helmink et al., 2020) (Fig. S1R). When the five datasets were analyzed together, we observed that the proportion of patients with tumor Treg accumulation after PD-1 monotherapy was significantly higher than the theoretical value of 50% corresponding to no increase (Fig. 1I). Thus, our meta-analysis of 53 metastatic melanoma patients with pre- and post-PD-1 immunotherapy biopsies suggests that the majority experienced Treg increase after treatment. Consequently, we set out to understand the relevance and mechanistic underpinnings of αPD-1-mediated Treg expansion in melanoma.

### αPD-1-mediated increase in tumor Treg numbers limits the effectiveness of immunotherapy

While previous reports showed that PD-1 immunotherapy synergizes with extensive Treg depletion (Arce Vargas et al., 2017; Dodagatta-Marri et al., 2019; Kidani et al., 2022), these studies do not establish a causal link between the αPD-1-mediated Treg increase and the outcome of immunotherapy. To investigate this question, we administered αPD-1 to D4M-S melanoma-bearing Foxp3^DTR^ mice, decreased Treg numbers to pre-therapy levels using a suboptimal dose of diphtheria toxin (DT), and measured tumor weight (Fig. 2A). Care was taken not to deplete Tregs completely. As expected, PD-1 inhibition increased tumor Tregs compared to isotype-treated mice. Co-administration of DT and αPD-1 reduced Treg percentages to the level of the isotype group in 13/27 (Treg Low) mice and was ineffective on the remaining mice (Treg Hi) (Fig. 2B, S2A-B). After αPD-1 treatment, 30% of mice had tumors of equivalent weight to those growing in animals receiving isotype control antibodies, while the remaining mice controlled tumor growth (Fig. 2C). Thus, our model captures the variable response patients display to PD-1 immunotherapy. Importantly, we observed that 43% of Treg Hi mice experienced uncontrolled tumor growth compared to 8% in the Treg Low group (Fig. 2C). These data indicate that the increase in Treg numbers due to PD-1 inhibition hinders tumor rejection.

**Figure 2.**
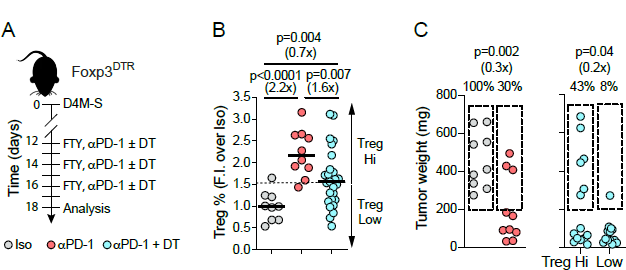
αPD-1-mediated Treg increase hinders tumor rejection. A. Experimental scheme for partial Treg ablation. **B.** Quantification of tumor Treg percentage fold increase over the isotype group. The dotted line indicates the threshold above which we consider Tregs increased (the mean between the highest isotype and lowest αPD-1 sample values). n=9 (Iso) 10 (αPD-1) and 27 (αPD-1 + DT) mice/group pooled from 2 separate experiments. Bars depict the median value of the distribution. p values by Mann-Whitney U test. **C.** Tumor weight of isotype and αPD-1 treated mice (left) and αPD-1 + DT treated mice (right). Mice treated with αPD-1 + DT are stratified by Treg Hi (n=14) and Low (n=13). Dotted rectangles indicate the tumors of 200 - 750 mg, corresponding to the tumor weight range in isotype-treated animals. p values by chi-squared test.

### αPD-1 does not increase TCR signaling in tumor Tregs

We hypothesized that tumor Treg accumulation following PD-1 blockade could be due to enhanced activation through the T cell receptor (TCR). Because TCR signaling is a significant inducer of Ca^2+^ influx in T cells, we monitored the levels of cytosolic Ca^2+^ ions using the genetically encoded indicator Salsa6f (Dong et al., 2017). Salsa6f is a fusion of tdTomato and GCaMP6f. TdTomato emits constant red fluorescence, whereas green fluorescence from GCaMP6f is proportional to the cytosolic concentration of Ca^2+^. We bred Foxp3^creERT2^ x Rosa26^LSL-Salsa6f^ mice that express Salsa6f stably and specifically in endogenous Tregs upon tamoxifen administration (Fig. 3A). To quantify tumor Treg activation in vivo, we implanted a D4M-S tumor in tamoxifen-treated Foxp3^creERT2^ x Rosa26^LSL-Salsa6f^ mice. Once each tumor was established, we installed a dorsal skinfold chamber (DSFC) to enable intravital imaging and performed functional intravital microscopy (F-IVM) (Fig. 3B). We visualized Tregs at the tumor-stroma border, as we previously described (Marangoni et al., 2021), and could readily distinguish resting and activated states based on Salsa6f red and green fluorescence signals (Fig. S3A and Supplemental Movie 1). We observed frequent Treg activation in both control mice and mice treated with αPD-1 24h earlier (Fig. 3C and Supplemental Movie 2). For each cell track, we quantified GFP intensity over time and subtracted the baseline signal (see Methods). We identified several peaks of GFP fluorescence, corresponding to individual instances of activation (Fig. 3D). Approximately 30% of endogenous Tregs in both groups signaled during the observation window (Fig. 3E), in line with previous findings using adoptively transferred NFAT-GFP expressing Tregs in a different tumor model (Marangoni et al., 2021). To accurately quantify activation, we focused on track segments corresponding to signaling peaks (Fig. 3F). The percentage of time an individual Treg was observed signaling was equivalent in control and αPD-1 treated mice (Fig. 3G). The maximum fluorescence increased while signaling duration decreased in the αPD-1 group, resulting in a comparable area under the curve (AUC) between control and αPD-1 treated mice (Fig. 3H).

**Figure 3.**
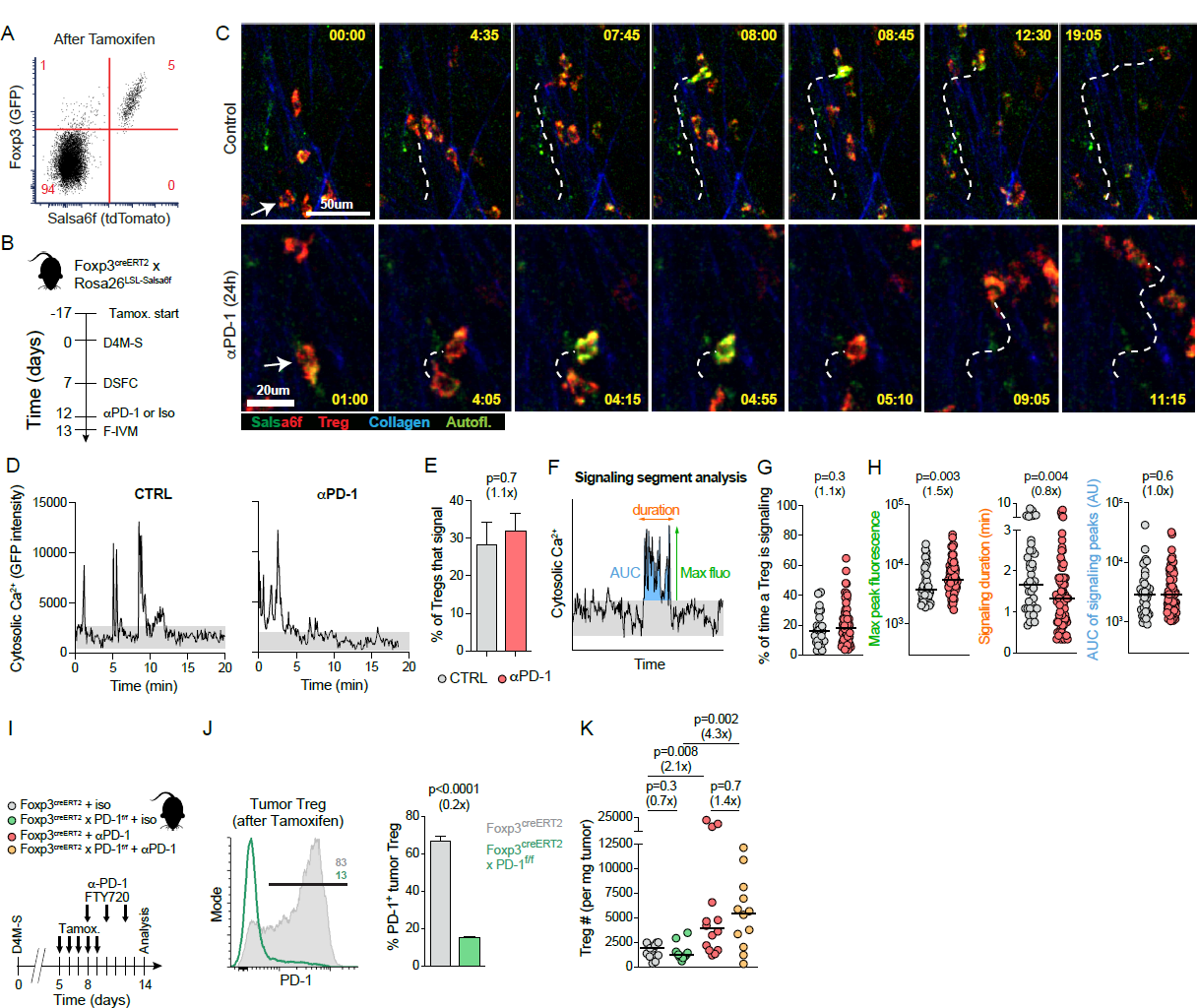
Indirect mechanisms are the main driver of tumor Treg accumulation after PD-1 blockade. A. Representative dot plot showing Salsa6f expression in blood Tregs from one representative tamoxifen-treated Foxp3^creERT2^ x Rosa26^LSL-Salsa6f^ mouse. **B.** Experimental scheme for F-IVM. **C.** Image sequences illustrating tumor Treg motility and Ca^2+^ signaling reported by Salsa6f in control or αPD-1 treated mice. Arrows point to the tracked cell, and the dotted line illustrates the cell trajectory. Time in min: sec. **D.** Dynamics of GFP fluorescence intensity signals in representative Treg tracks. Track-specific baseline is depicted in grey. **E.** Percentage of Treg tracks displaying at least one signaling peak. Mean ± SEM is shown. p values by Student’s *t*-test. **F.** Illustration of track segments and associated parameters. **G.** Percentage of time a Treg is signaling. **H.** Quantification of maximum GFP fluorescence, signaling duration, and area under the curve (AUC) for individual signaling segments. In **G and H**, bars represent medians. p values by Mann Whitney U test. For **C-H**, we analyzed 5 control and 7 αPD-1 movies, corresponding to 115 control and 207 αPD-1 Treg tracks and 41 control and 86 αPD-1 signaling segments. **I.** Scheme of Treg-specific PD-1 deletion experiments. **J.** Representative histogram and quantification of PD-1 expression in tumor Tregs within Foxp3^creERT2^ or Foxp3^creERT2^ x PD-1^f/f^ mice treated with tamoxifen. Mean ± SEM is depicted. p values by Student’s *t*-test. **K.** Treg counts per mg of tumor in Foxp3^creERT2^ or Foxp3^creERT2^ x PD-1^f/f^ mice treated with αPD-1 or isotype control antibodies. n=12 to 14 per group from three independent experiments. Bars represent medians. p values by Mann Whitney U test.

To investigate whether these slight variations in signaling dynamics imposed by αPD-1 affect Treg accumulation, we generated Foxp3^creERT2^ x PD-1^f/f^ mice to selectively delete PD-1 on Tregs by administration of tamoxifen (Fig. 3I). The efficiency of PD-1 deletion was ∼80% (Fig. 3J). Even when excluding CD44^lo^ Tregs that were Ki67 negative and may not respond to antigens in the tumor environment, we observed only a small but significant increase in Ki67 expression in PD-1-deleted compared to PD-1-sufficient Tregs in the same mouse. GITR or ICOS remained unchanged (Fig. S3B). Importantly, Treg-specific PD-1 deletion alone did not increase tumor Treg numbers compared to control Foxp3^creERT2^ mice (Fig. 3K), demonstrating that the moderate influence of PD-1 blockade on TCR-mediated Treg activation and Ki67 expression is insufficient to trigger intratumor accumulation. In stark contrast, antibody-mediated PD-1 blockade increased Treg numbers irrespective of their PD-1 expression (Fig. 3K), revealing that αPD-1 causes the numerical expansion of tumor Tregs through a cell-extrinsic mechanism.

### CD8 cells and IL-2 are required for αPD-1-mediated tumor Treg accumulation

Considering the pronounced response of CD8 cells within αPD-1 treated D4M-S melanoma, we hypothesized that CD8 cells may support Treg accumulation during PD-1 blockade. By analyzing D4M-S tumors explanted from Foxp3^GFP^ x E8I^cre^ x Rosa26^LSL-Tomato^ mice that allow for visualization of CD8 and Tregs, we observed that these cells clustered together in several areas of the tumor (Fig 4A). To assess co-localization, we measured the distance between each Treg and the closest CD8 cell in the original dataset, and after randomization of Treg positions (Fig S4A). The median distance between Treg and CD8 cells in the original dataset was significantly lower (10μm) than the distance after Treg shuffling (15μm), demonstrating that there is a non-random distribution of Tregs relative to CD8 cells (Fig 4B). To test whether tumor Treg increase after αPD-1 depends on CD8 cells, we depleted CD8 cells during PD-1 therapy (Fig 4C). We observed an accumulation of Tregs with αPD-1 compared to isotype controls, which depended entirely on CD8 cells (Fig 4D). Conversely, αPD-1-mediated Treg accumulation in tumor-draining lymph nodes occurred irrespectively of CD8 cells (Fig S4B).

**Figure 4.**
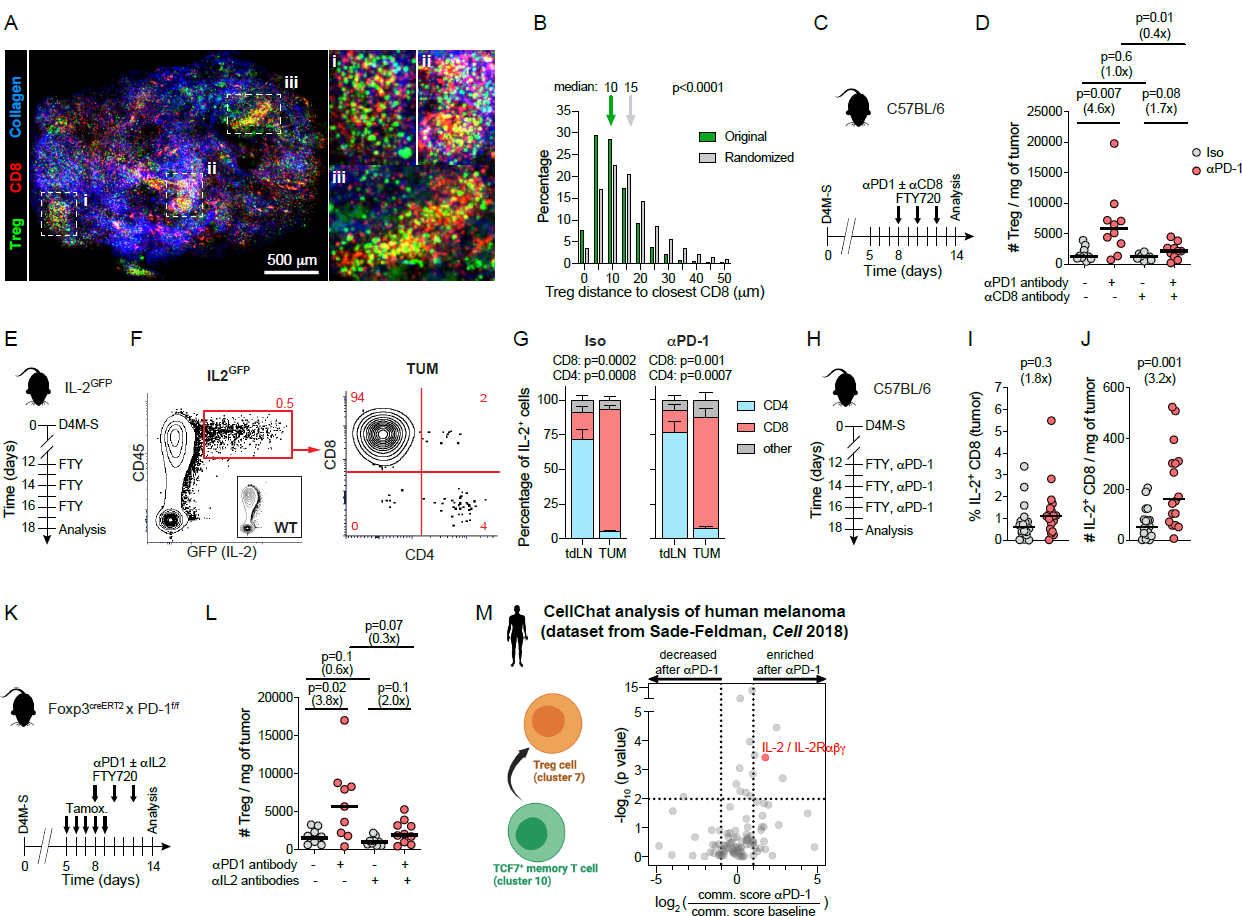
Tumor Treg accumulation after PD-1 blockade depends on CD8 cells and IL-2. A. Representative multiphoton montage image of a D4M-S tumor explanted from a Foxp3^GFP^ x E8I^cre^ x Rosa26^LSL-Tomato^ mouse. Three regions of interest are highlighted and magnified on the right. One tumor representative of three is shown. **B.** Distribution of Treg distance to closest CD8 cell in both the original image and after randomization of Treg positions. p values by Mann Whitney U test. **C.** Experimental scheme for CD8 depletion experiments. **D.** Treg numbers per mg of D4M-S tumor in mice treated or not with αPD-1 and CD8 depleting antibodies. n=10 mice/group pooled from 2 independent experiments. **E.** Experimental scheme to determine the source of IL-2 using IL-2^GFP^ mice. **F.** Representative dot plot of IL-2-transcribing cells (GFP^+^) in an IL2^GFP^ mouse. Inset shows a wild-type mouse as a negative control. By gating GFP^+^ cells, we determined if IL-2-transcribing cells expressed CD8^+^ or CD4^+^. **G.** Proportions of CD4 and CD8 cells among IL-2 producers in tumor-draining lymph nodes (tdLN) and D4M-S melanomas within IL2^GFP^ mice treated as indicated. Mean ± SEM of three independent experiments is depicted. p values by Student’s *t*-test. **H.** Experimental scheme for IL-2 protein quantification after PD-1 immunotherapy. **I, J.** Percentage (**I**) and counts (**J**) of IL-2-producing tumor CD8 cells with or without PD-1 blockade. n=18 mice per group in four independent experiments. **K.** Experimental scheme for IL-2 neutralization. **L.** Treg numbers per mg of D4M-S tumor in mice treated or not with αPD-1 and antibodies neutralizing the interaction of IL-2 with α and β subunits of the IL-2 receptor. n=9 to 10 mice/group pooled from 2 separate experiments. In **D, I, J, L,** bars depict medians and p values by Mann Whitney U test. **M.** CellChat analysis of communication pathways in human melanoma. *TCF7*-expressing memory T cells and Tregs correspond to clusters 10 and 7 of (Sade-Feldman et al., 2018). The volcano plot depicts the ratio of communication score after vs. before PD-1 immunotherapy, and the p value (Mann-Whitney test) of ligand upregulation following PD-1 blockade. The IL-2 / IL2Rαβγ pathway is highlighted in red. All other interactions which were identified by CellChat and which have a finite log-fold change of communication probability are shown in grey.

We set out to identify what factor expressed by activated tumor CD8 cells supports tumor Tregs after PD-1 blockade. We reasoned that the candidate molecules 1) should be produced mainly by CD8 cells in the tumor but not in lymph nodes since CD8 depletion only affected Treg expansion in the tumor (compare Fig. 4D to Fig. S4B), and 2) should prolong Treg survival due to limited evidence of enhanced proliferation in tumor Tregs after αPD-1 treatment (Fig. 1E, F). IL-2 was a prime candidate as it is a crucial trophic and survival factor for Tregs (Chinen et al., 2016; Owen et al., 2018; Smigiel et al., 2014), and a recent report characterized IL-2 production by *Tcf7*-expressing CD8 memory cells in the context of viral infection (Kahan et al., 2022). *Tcf7*-expressing CD8 cells infiltrate tumors and are exquisitely responsive to PD-1 immunotherapy (Siddiqui et al., 2019). We thus postulated that CD8 cells can be an intratumoral source of IL-2. To examine this possibility, we implanted D4M-S tumors into IL-2^GFP^ mice (Ditoro et al., 2018) to identify IL-2 transcribing cells by GFP expression (Fig 4E). In line with previous studies (Liu et al., 2015; Pierson et al., 2013), the main source of IL-2 in the lymph node was Th cells, while CD8 cells only accounted for ∼20% of IL-2-producing cells. In contrast, CD8 cells were the primary producer of IL-2 in the tumor (∼90% of IL-2-producing cells) independently of PD-1 blockade (Fig 4F, G). To study if αPD-1 treatment increases IL-2, we analyzed IL-2 protein production by tumor CD8 cells in response to αPD-1 in C57BL/6 mice (Fig 4H). We found no significant increase in the production of IL-2 on a per-cell basis (Fig. 4I, S4C); however, there was an expansion of CD8 cells per mg of tumor after PD-1 blockade and therefore increased numbers of IL-2-producing CD8 cells (Fig 4J). To investigate if IL-2 is required for tumor Treg accumulation in response to αPD-1, we blocked IL-2 binding to the α and β subunits of its receptor through neutralizing antibodies (Fig 4K). Comparable to our results for CD8 depletion, αPD-1 triggered an increase in Treg numbers that was entirely abrogated when αPD-1 was given simultaneously with IL-2 neutralizing antibodies (Fig 4L). αPD-1-mediated Treg accumulation in the lymph node was only partially dependent on IL-2 (Fig. S4D), consistent with the notion that IL-2 supports cTreg but not eTreg homeostasis (Smigiel et al., 2014). Together, these data strongly suggest that tumor Treg accrual triggered by αPD-1 depends on CD8-produced, intratumoral IL-2.

### Upregulation of the CD8 / IL-2 / Treg axis in human melanoma patients receiving PD-1 immunotherapy

To assess whether the CD8 / IL-2 / Treg axis we identified in mice is also present in human melanomas, we performed CellChat analysis (Jin et al., 2021) on a published single-cell RNA sequencing dataset (Sade-Feldman et al., 2018). CellChat uses an accurately annotated receptor-ligand library to calculate the “interaction strength” between all cell clusters in a single-cell RNAseq experiment. To eliminate possible confounding factors represented by the variable treatment of patients, we included only cases treated with αPD-1 monotherapy; the control group comprised all the tumors sampled before treatment. We found that PD-1 immunotherapy upregulated IL-2 production in the cluster marked by *TCF7* expression, a gene mainly expressed by CD8 T cells in the tumor (Sade-Feldman et al., 2018). CellChat predicted that PD-1 immunotherapy increases the IL-2 / IL-2Rαβγ pathway of communication between *TCF7*-expressing memory T cells and Tregs in human melanoma (Fig. 4M). Thus, CellChat predicts that PD-1 blockade triggers a CD8 / IL-2 / Treg axis in melanoma patients, suggesting that our mechanistic studies in mice have human relevance.

### Accumulation of tumor Tregs following αPD-1 treatment depends on the TCR and CD28

Some in vitro studies propose that IL-2 and CD28 may drive Treg expansion in the absence of TCR-mediated signaling (He et al., 2017; Li et al., 2012). If TCR-independent tumor Treg accumulation occurred after PD-1 blockade, bystander Tregs with no specificity for tumor antigens might participate in local immunosuppression with tumor antigen-specific Tregs. We investigated this possibility by assessing tumor Treg levels in mice with Treg-specific, inducible deletion of either TCRα or CD28 (Foxp^creERT2^ x TCRα^f/f^ or Foxp3^creERT2^ x CD28^f/f^). These mice received D4M-S tumors, tamoxifen, and αPD-1 before Treg analysis (Fig S4E). Tumor Tregs in Foxp^creERT2^ x TCRα^f/f^ mice exhibited 70% TCR deletion, while Foxp3^creERT2^ x CD28^f/f^ mice showed 55% CD28 deletion (Fig S4F, G). Because TCR or CD28 deletion was not complete, we performed subsequent analyses by gating on Tregs that were negative for these genes. While PD1 blockade in Foxp3^creERT2^ mice increased the levels of tumor Tregs two-fold compared to isotype controls, Tregs negative for the TCR or CD28 failed to expand (Fig S4H). Together, these data demonstrate that TCR and CD28 signaling is required for the αPD-1-driven, CD8 and IL-2-dependent expansion of Treg cells.

### IL-2-mediated tumor Treg accumulation depends on ICOS

To study the effect of IL-2 on tumor Tregs, we administered IL-2 immunocomplexes (IL-2i.c.) that direct the effects of IL-2 to cells expressing IL-2Rα (Boyman et al., 2006) (Fig 5A). We performed these experiments in D4M melanoma because it contains a small amount of Tregs, and thus the effects of IL-2i.c. may be the most evident. Indeed, we observed a marked increase in tumor Treg numbers in response to αPD-1 and IL-2i.c., compared to αPD-1 only (Fig 5B). This accumulation of tumor Tregs was not caused by increased proliferation (Fig 5C) or Treg influx from the lymph node (blocked by FTY720), suggesting enhanced survival. In addition, the expression of pro-apoptotic Bim increased after IL-2i.c. administration (Fig. 5D), leading us to consider the IL-2-dependent anti-apoptotic factors Bcl-2, Bcl-xL, and Mcl-1 as possible candidates mediating prolonged tumor Treg survival (Pierson et al., 2013; Wang et al., 2012; Whitehouse et al., 2017). This mechanism would be reminiscent of lymph node cTregs, where high amounts of Bim appear to be balanced by high expression of Bcl-2 (Fig. S5). However, Bcl-2, Bcl-xL, and Mcl-1 were not upregulated in tumor Tregs after treatment with αPD-1 and IL-2i.c. compared to αPD-1 alone (Fig 5E). IL-2i.c. treatment instead increased the expression of the costimulatory and anti-apoptotic molecule ICOS (Fig 5F). We thus blocked the interaction between ICOS and its only ligand (ICOSL) through αICOSL antibodies, which have been previously validated (Smigiel et al., 2014). ICOSL blockade prevented IL-2-mediated tumor Treg accumulation (Fig 5G, H), demonstrating a critical role for the IL-2 / ICOS axis in orchestrating Treg abundance in melanoma. To assess whether ICOSL blockade may specifically target Tregs, we measured ICOS expression on T cells after treatment with αPD-1 antibodies compared to isotype controls (Fig. 5I). We found that ICOS is expressed at the highest levels on tumor Tregs, followed by Th and CD8 cells. ICOS expression was further increased by PD-1 blockade, especially on tumor Tregs (Fig 5J).

**Figure 5.**
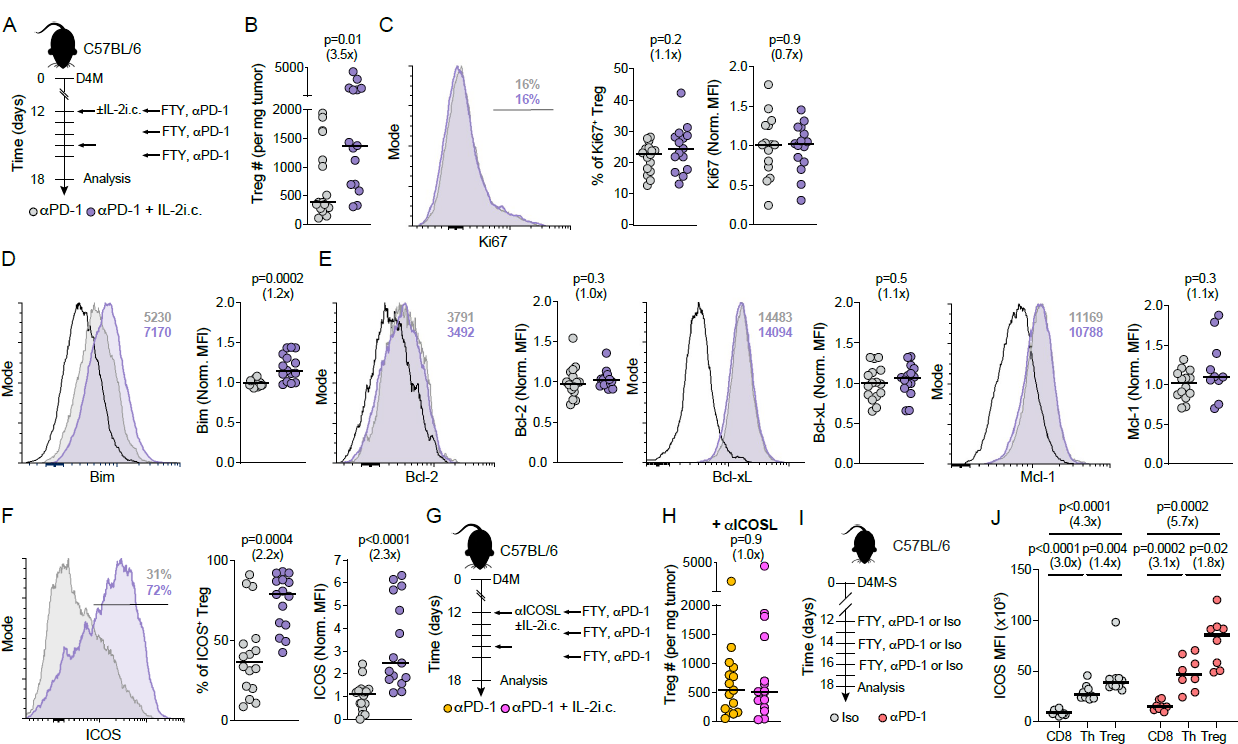
ICOS mediates IL-2-driven tumor Treg accumulation. A. Experimental scheme for stimulation of tumor Tregs with IL-2 immunocomplexes (i.c.). **B.** Treg number per mg of D4M melanoma treated with αPD-1 ± IL-2i.c. **C.** Representative histogram and quantification of Ki67 expression in tumor Tregs treated with PD-1 blockade with or without IL-2i.c. **D, E.** Representative histograms and normalized MFI of the pro-apoptotic marker Bim (**D**) and the anti-apoptotic molecules Bcl-2, Bcl-xL, and Mcl-1 (**E**) in tumor Tregs after administration of αPD-1 with or without IL-2i.c. **F.** Representative histogram and quantification of ICOS expression in tumor Tregs treated with PD-1 blockade ± IL-2i.c. **G.** Experimental scheme for ICOSL blockade. **H.** Tumor Treg numbers upon treatment with αPD-1 and αICOSL, with or without IL-2i.c. For **B-H**, n=10-16 mice/group pooled from 2-3 separate experiments. **I.** Experimental scheme for measuring ICOS expression on T cells during PD-1 blockade. **J.** MFI of ICOS on tumor-associated CD8, Th, and Treg cells treated with isotype or αPD-1. n=8-10 mice/group from 2 separate experiments. p values by Mann Whitney U test.

### Concurrent ICOSL / PD-1 blockade does not improve the effectiveness of PD-1 immunotherapy

Since ICOSL blockade inhibited the CD8- and IL-2-mediated support to Tregs triggered by αPD-1, it may synergize with PD-1 blockade to increase melanoma rejection. On the other hand, blockade of ICOS signaling on effector CD8 and Th cells may decrease their anti-tumor potential. To distinguish between these possibilities, we treated C57BL/6 mice bearing immunogenic or non-immunogenic melanomas with concomitant ICOSL / PD-1 blockade and measured tumor growth (Fig 6A and S6A). The non-immunogenic D4M melanoma was insensitive to PD-1 blockade, consistent with previous reports (Lo et al., 2021), and combination treatment with αICOSL did not modify this outcome (Fig S6A). In contrast, αPD-1-treated mice better controlled immunogenic D4M-S tumors than isotype-treated mice. αICOSL alone did not affect D4M-S growth and, when administered in combination with αPD-1, did not improve the efficacy of αPD-1 monotherapy (Fig 6B). We thus hypothesized that the favorable effects of ICOSL blockade on tumor Tregs are counterbalanced by a detrimental impact on antitumor immunity.

**Figure 6.**
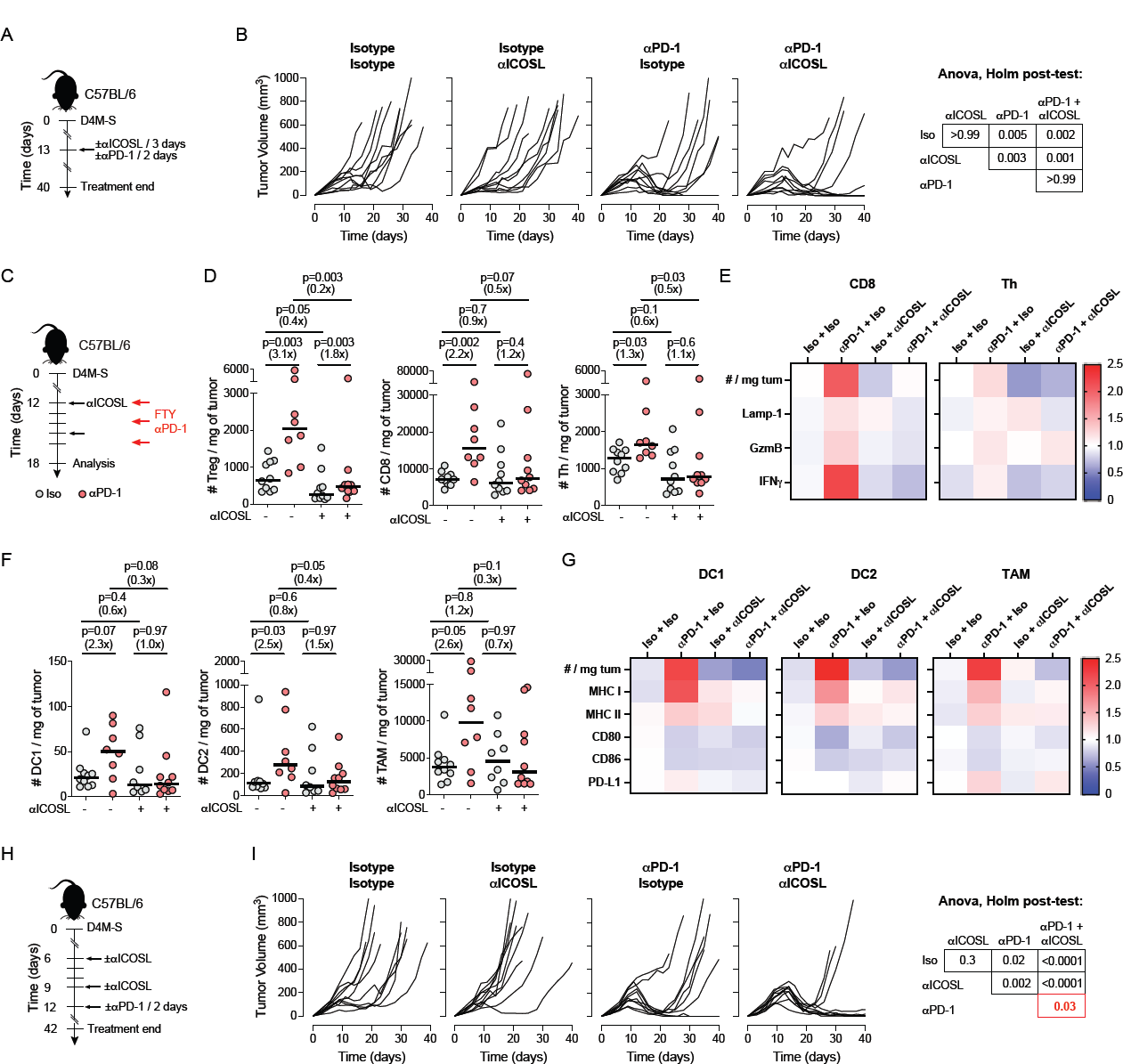
Effectiveness of sequential, but not concomitant, ICOSL / PD-1 immunotherapy. A. Experimental scheme for concomitant ICOSL / PD-1 therapy. **B.** Growth curves of D4M-S tumors in wild-type mice treated with αICOSL or αPD-1 antibodies individually or in combination. Each line represents one mouse. n=10 mice/group from 2 separate experiments. p values by type II Anova followed by Holm post-test. **C.** Experimental scheme to characterize the effects of ICOSL / PD-1 therapy on T cells and APCs. **D.** Treg, CD8, and Th cell numbers per mg of D4M-S melanoma treated with αPD-1 or αICOSL antibodies individually or in combination. **E.** Heatmap depicting cell numbers per mg of D4M-S melanoma, Lamp-1 and Granzyme B expression, and number of IFNγ-producing cells per mg of tumor after administration of the indicated treatments. Colors represent the median of all experimental values, normalized by the average of the iso+iso group. **F.** DC1, DC2, and TAM cell numbers per mg of D4M-S melanoma treated with αPD-1 or αICOSL antibodies alone or in combination. **G.** Heatmap depicting cell numbers per mg of D4M-S melanoma and MHCI, MHCII, CD80, CD86, and PD-L1 expression in tumor DC1s, DC2s, and TAMs after administration of the indicated treatments. Colors represent the median of all experimental values, normalized by the average of the iso+iso group. For **D-G,** n=8-10 mice/group pooled from 2 separate experiments. Bars depict medians. p values by Mann-Whitney U test. **H.** Experimental scheme for sequential ICOSL / PD-1 blockade. **I.** Growth curves of D4M-S tumors in wild-type mice treated with monotherapies or sequential αICOSL or αPD-1 immunotherapy. Each line represents one mouse. n=10 mice/group pooled from 2 separate experiments. p values by type II Anova followed by Holm post-test.

### Impact of ICOSL blockade on the antitumor immune response

We characterized the effects of αICOSL alone or in combination with αPD-1 on Treg, CD8, and Th cells (Fig 6C). ICOSL monotherapy decreased Treg cells more profoundly than Th and CD8 cells (Fig 6D). As a result, we observed a higher CD8/Treg ratio in mice treated with ICOSL blockade (Fig S6B), which predicts a good prognosis in human cancer patients (Sato et al., 2005). PD-1 monotherapy increased the numbers of Treg and CD8 cells and, in this round of experiments, weakly increased Th cell counts as well. However, when ICOSL and PD-1 blockade were combined, the αPD-1-mediated increase in Treg, Th, and CD8 cell numbers was negated (Fig 6D). In tumor-associated CD8 cells, the expression of granzyme B but not LAMP-1 decreased significantly upon blockade of ICOSL and PD-1 compared to αPD-1 monotherapy. Additionally, there was a tendency toward reduced numbers of IFNγ-producing cells. (Fig. 6E, S6C). We observed a similar trend in Th cells, even though PD-1 blockade enhanced their effector functions to a lower level than CD8 cells (Fig. 6E, S6D). We also assessed the impact of ICOSL / PD-1 therapy on tumor-associated antigen-presenting cells (APCs), because they express ICOSL (Aicher et al., 2000). We identified XCR1-expressing DC1, Sirpα-expressing DC2, and tumor-associated macrophages (TAMs) as shown in Fig S6E. Similarly to what we observed for effector T cells, the combination of αICOSL and αPD-1 antibodies resulted in less tumor DC1, DC2, and TAMs compared to PD-1 blockade alone (Fig 6F, G). Additionally, there was a trend toward impaired antigen presentation (MHC-I) with no change in MHC-II, CD80, CD86, and PD-L1 expression on DC1, DC2, and TAMs during ICOSL / PD-1 blockade compared to PD-1 monotherapy (Fig 6G, Fig S6F-I). αICOSL monotherapy did not change these parameters. These data indicate that while αICOSL alone impairs Tregs with negligible effects on other tumor-associated immune cells, its combination with αPD-1 prevents the enhancement in effector T cell and APC numbers and functions normally triggered by PD-1 monotherapy.

### Sequential ICOSL / PD-1 blockade improves the effectiveness of PD-1 immunotherapy

We explored whether we could take advantage of the benefits of ICOSL blockade while avoiding its detrimental effects upon αPD-1 co-administration. We reasoned that, by first administering αICOSL antibodies to D4M-S bearing mice, we could preferentially decrease tumor Treg numbers and enhance the intratumor CD8 / Treg ratio so that subsequent PD-1 blockade (Fig 6H) would improve the antitumor efficacy of CD8 cells. We observed no benefit from αICOSL treatment compared to mice receiving isotype-matched antibodies, while PD-1 monotherapy resulted in better tumor control (Fig 6I). Importantly, sequential ICOSL / PD-1 blockade significantly improved tumor control over PD-1 monotherapy (Fig 6I). Thus, we provide evidence of synergy between sequentially administered ICOSL and PD-1 blockade against immunogenic melanoma.

## DISCUSSION

In this study, we found that PD-1 blockade promotes the crosstalk between tumor-associated CD8 and Treg cells, increasing tumor Treg numbers and reducing the efficacy of immunotherapy. We used genetic deletion and multiphoton intravital microscopy to show that interrupting cell-intrinsic PD-1 signaling has a limited impact on TCR-mediated activation and Treg accumulation within the tumor. Instead, αPD-1-mediated tumor Treg increase was dependent on CD8-provided IL-2 and was mediated by ICOS expression on Tregs. We then targeted ICOS signaling to block the CD8:Treg crosstalk and improve the efficacy of PD-1 immunotherapy. We found that concomitant PD-1 and ICOSL blockade did not enhance the effectiveness of PD-1 monotherapy since αICOSL counteracted αPD-1-mediated APC and T cell effector functions. However, αICOSL and αPD-1 synergized when administered sequentially.

A seminal report showed that PD-1 blockade increases the abundance and activation of tumor-associated Tregs in patients with hyper-progressive gastro-esophageal cancer (Kamada et al., 2019). However, there is no consensus on the effect of PD-1 blockade on melanoma-associated Tregs: one study detected no differences in Treg abundance after treatment (Ribas et al., 2016), while a subsequent report showed Treg enrichment in the blood of patients not responding to PD-1 immunotherapy (Woods et al., 2018). We addressed these discrepancies by performing a meta-analysis on data from patients treated with PD-1 monotherapy only, and for whom paired biopsies taken before and on-treatment were available. We found that PD-1 immunotherapy increased the number of tumor Tregs in the majority of patients. While melanoma is generally immunogenic, variation among individuals is high (1-100 mutations per Mb of exome) (Lawrence et al., 2013). Therefore, patients who did not exhibit an increase in Tregs following αPD-1 treatment may have had a poorly immunogenic cutaneous melanoma, or a subtype not caused by ultraviolet exposure (e.g., uveal or mucosal). Our study also suggests that early reports may have been confounded by the variable abundance of Tregs in individual patients at baseline and by the inclusion of patients undergoing disparate treatments, often including CTLA-4 blockade.

A comparison of our current and published data (Marangoni et al., 2021) indicates that PD-1 and CTLA-4 blockade impact tumor Treg activation through different mechanisms. While PD-1 blockade induced both tumor CD8 and Treg cell expansion, with a state of Treg activation characterized by low proliferation and high expression of Foxp3, GITR, and ICOS, CTLA-4 inhibition resulted in tumor Treg proliferation and accumulation with no enhancement in activation markers (Marangoni et al., 2021). These differences may be explained by the ability of PD-1 to modulate TCR and CD28 signaling (Hui et al., 2017; Yokosuka et al., 2012) while CTLA-4 primarily modulates CD28 signaling (Qureshi et al., 2011), or by the distinct effects of each immunotherapy on tumor-associated CD8 and Th cells (Wei et al., 2017). We also analyzed the impact of PD-1 blockade on lymph node Tregs. We observed a trend towards increased eTreg percentage in tumor-draining lymph nodes after αPD-1 administration, and these eTregs were more proliferative. CD44^hi^CD62L^−^ CD8 percentage was also significantly enhanced. All these findings agree with a previous study on transgenic mice with T cell-specific PD-1 deficiency (Kamada et al., 2019). However, we noticed that in response to αPD-1, the characteristics of Treg activation in lymph nodes and the tumor environment differed: lymph node Tregs display more prominent proliferation but no upregulation of activation markers. The reason for this phenomenon could be that Tregs adapt to different tissues by taking on specific transcription signatures (Panduro et al., 2016). In particular, lymph node eTreg cells must complete several differentiation steps to become Treg residing in non-lymphoid tissues, including tumors (Miragaia et al., 2019). The notion that the Treg transcriptome dynamically changes during their development fits with our observation that PD-1 signaling is particularly important during the cTreg to eTreg transition but less so in later phases of differentiation.

Our intravital imaging studies on tumor Treg activation showed that αPD-1 antibodies did not increase TCR signaling, and Treg-specific genetic deletion of PD-1 did not cause their accumulation in immunogenic melanoma. These observations appear to conflict with published reports on pancreatitis (Zhang et al., 2016), experimental autoimmune encephalomyelitis and type 1 diabetes (Tan et al., 2021), *Toxoplasma gondii* infection (Perry et al., 2022), and tumors (Kumagai et al., 2020) that indicated a cell-intrinsic control of Treg numbers and activation by PD-1. However, indirect effects may be dominant over cell-intrinsic consequences of PD-1 blockade when it comes to controlling Treg numbers in immunogenic tumors. In support of this idea, we did observe increased expression of Ki67 in PD-1-deleted compared to PD-1-expressing Tregs in the same mice, yet this direct effect was insufficient to cause tumor Treg accumulation. While we cannot exclude that the cell-intrinsic PD-1 deletion on Tregs increases their suppressive function, we demonstrate that Treg number increase, an indirect effect of PD-1 blockade, significantly contributes to tumor growth (Fig. 2). We speculate this is not the only mechanism responsible for Treg accumulation after PD-1 immunotherapy. Tumor Tregs respond to self-antigens (Malchow et al., 2013) as well as some mutational neo-antigens (Ahmadzadeh et al., 2019), so if a tumor expresses self-antigens ectopically or in high amounts (Coulie et al., 2014) or Treg-specific neoantigens, PD-1 blockade might directly enhance local Treg proliferation. The reason why tumor Tregs, which express PD-1 (Kamada et al., 2019), did not show increased TCR signaling after PD-1 blockade remains to be determined. Since tumor Tregs are extensively activated through the TCR even before the administration of immunotherapy ((Marangoni et al., 2021) and Fig. 3E), the direct effects of PD-1 blockade on TCR-mediated activation may be minor.

One key finding of our study is that CD8 cells colocalize with Tregs in tumors and mediate αPD-1-dependent Treg increase. Tumor-associated CD8 cells were the dominant source of intratumor IL-2, a surprising finding since Th cells are the primary producers of IL-2 in secondary lymphoid organs (Boyman and Sprent, 2012). In addition, it has been challenging to analyze IL-2 production within the tumor microenvironment and in the absence of ex vivo restimulation, at least in part because of limited assay sensitivity (Selby et al., 2013) (John Engelhardt, personal communication). One study on sorted tumor T cell populations found that Th and CD8 cells upregulated the IL-2 transcript after CTLA-4 immunotherapy (Hannani et al., 2015). Our recent investigation using IL-2^GFP^ mice confirmed that tumor-associated Th and CD8 cells produce similar levels of IL-2 per cell (Marangoni et al., 2021). Because Th cells are also boosted by PD-1 immunotherapy (Nagasaki et al., 2020), the primary source of IL-2 is likely determined by the relative abundance of CD8 and Th cells within the tumor environment. IL-2 is preferentially secreted at the immunological synapse between effector T cells and APCs (Huse et al., 2006) but eventually diffuses within tissues (Liu et al., 2015; Oyler-Yaniv et al., 2017). Resolution of the immunological synapse is a likely mechanism by which IL-2 is released into the intercellular space; if so, the instability of immune synapses between tumor-associated CD8 and tumor cells (Marangoni et al., 2013) might contribute to IL-2 dissemination within the tumor environment. We showed tumor Treg cells interpret IL-2 signaling by upregulating the co-stimulatory and antiapoptotic molecule ICOS. Previous work defined IL-2 and ICOS as necessary for maintaining cTreg and eTreg homeostasis, respectively, in secondary lymphoid organs (Smigiel et al., 2014). Our data extend these findings by demonstrating that IL-2 induces ICOS in tumor Tregs.

We found that αICOSL did not synergize with simultaneous PD-1 immunotherapy because it impacted not only Tregs but also αPD-1-stimulated effector T cells. The observation that CD8 cells were the primary source of IL-2, irrespective of PD-1 blockade, opened the possibility that the CD8 / IL-2 / Treg axis could be already active before immunotherapy, albeit at lower levels. We therefore pre-treated melanoma-bearing mice with αICOSL to reduce Tregs via inhibition of the CD8:Treg crosstalk, and subsequently administered PD-1 immunotherapy to boost CD8 activation, which led to synergy between treatments. Thus, our studies support the emerging concept that immunotherapies targeting Treg but having some impact on effector T cells should be administered to condition the immune environment before switching to a second therapeutic intervention to enhance CD8 stimulation (Ha et al., 2019). In addition, ICOSL / PD-1 blockade may be advantageous in settings of neoadjuvant immunotherapy, which is emerging as an exciting strategy to treat patients at high risk of developing metastatic disease (Patel et al., 2023). Indeed, a recent paper demonstrated that Treg inhibition during neoadjuvant immunotherapy increases the survival of mice bearing metastatic mammary tumors (Blomberg et al., 2023).

Our finding that ICOSL blockade boosts PD-1 immunotherapy contrasts with the current notion that ICOS should be stimulated, rather than blocked, to increase effector T cell functions and promote tumor rejection. However, this concept was developed in the context of CTLA-4 immunotherapy, which specifically induces a population of ICOS-expressing Th cells (Wei et al., 2017) that are the target for ICOS agonism (Chen et al., 2014; Fu et al., 2011). Therefore, the pattern of ICOS expression induced by distinct immunotherapies may determine whether ICOS signaling should be triggered or blocked to improve antitumor efficacy.

Human αICOSL antibodies are in clinical development to treat autoimmune diseases (Cheng et al., 2018; Sullivan et al., 2016), and they may be repurposed to treat immunogenic cancers in asynchronous combination with αPD-1. ICOSL blockade offers a better safety profile than ICOS-depleting antibodies, which likely eliminate ICOS-expressing Tregs in tumors and other non-lymphoid tissues, increasing the risk of autoimmunity. Since some patients treated with PD-1 monotherapy develop life-threatening immune-related adverse events (Wang et al., 2018), increasing the efficacy of immunotherapy without Treg depletion is paramount. The concept of interrupting CD8 cell-mediated and IL-2-dependent support to Tregs within the tumor environment might aid in achieving this goal.

## AUTHOR CONTRIBUTIONS

S.N.G. performed experiments, analyzed data, and wrote the paper; A.M. and R.S. performed experiments; R.S.S., B.L.W., and Q.N. did the bioinformatic analysis of human data; G.G. and M.H. tracked Treg cells in intravital microscopy experiments; T.R.M. provided some data on MC38 tumors; S.O. conducted intravital microscopy experiments and supervised movie analysis; F.M. conceived the project, performed experiments, analyzed data, and wrote the paper.

## ACKNOWLEDGMENTS

This work is funded by a pilot grant from the Chao Family Comprehensive Cancer Center of the University of California Irvine, a pilot grant from the Cancer Systems Biology Center of the University of California Irvine, and by the Melanoma Research Alliance – Bristol Myers Squibb Young Investigator Award #929155 (to F.M.), NSF grant DMS1763272 (to Q.N.), a grant from the Simons Foundation (594598, to Q.N.), and NIH grant R01AI168063 (to S.O.). A.M. is supported by T32 AR080622, and R.S.S by an NSF-GRFP fellowship. The authors thank Drs. Casey Weaver, Shimon Sakaguchi, Alexander Rudensky, and Michael Cahalan for providing transgenic mice, as well as Dr. Jennifer Wargo for sharing a bulk RNA sequencing dataset (Helmink et al., 2020).

---

**Figure S1.**
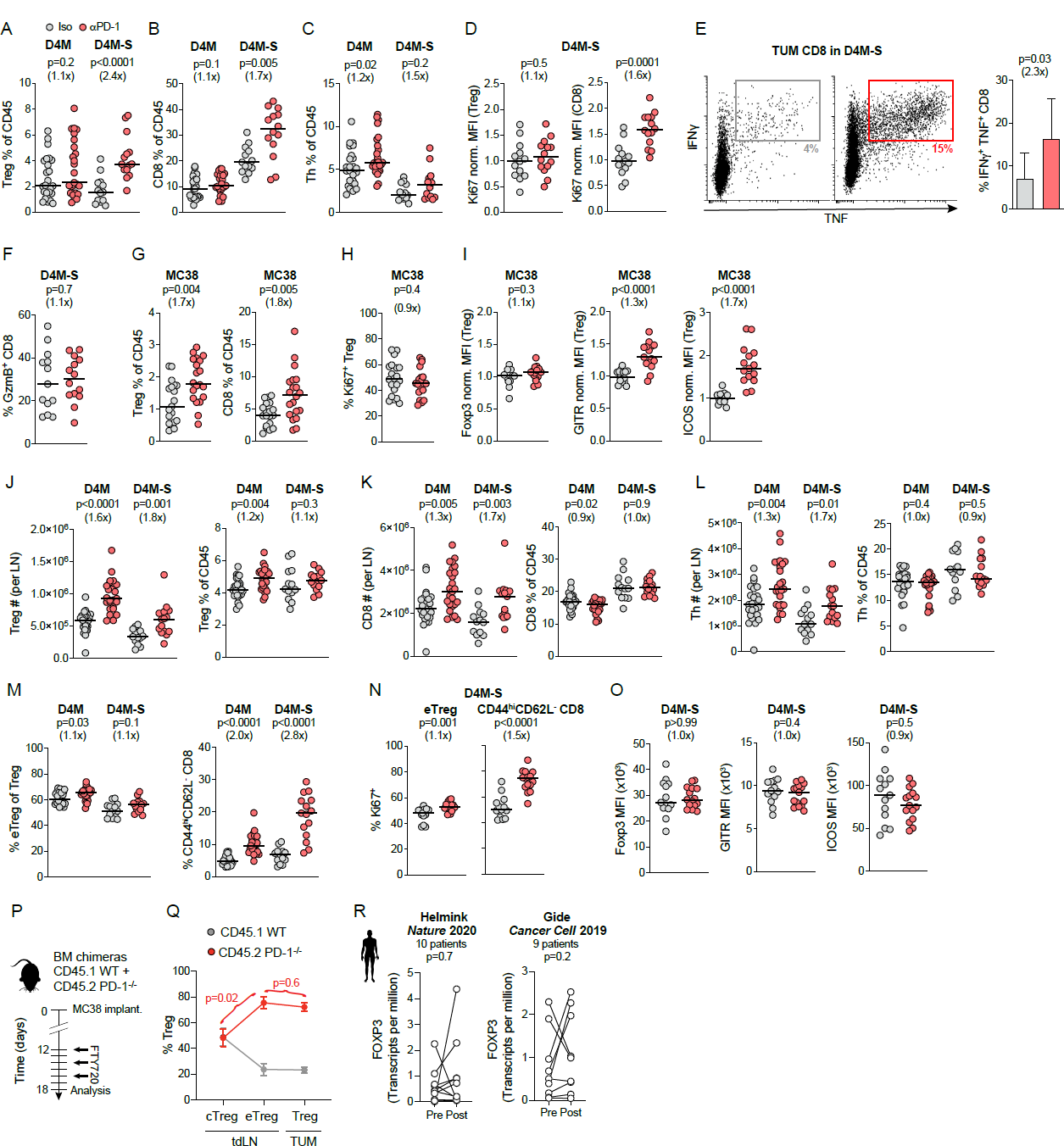
Effects of PD-1 immunotherapy differ from the lymph node to the tumor, related to Figure 1. **A-C.** Treg, CD8, and Th cells as a percentage of CD45^+^ in D4M or D4M-S tumors. **D.** Normalized MFI of Ki67 in Treg and CD8 cells from D4M-S tumors. **E.** Representative flow plots and quantification of IFNγ and TNF production by tumor CD8 cells restimulated for 8h with αCD3/αCD28 antibodies in the presence of brefeldin A. **F.** Granzyme B expression in CD8 cells from D4M-S tumors. **G.** Treg and CD8 cell percentage in MC38 tumors. **H-I.** Percentage of Ki67 expressing cells (**H**), and normalized Foxp3, GITR, and ICOS MFI (**I**) in Tregs from MC38 tumors. For **A-D** and **F-I,** n=25 (D4M), 12-15 (D4M-S), and 15-20 (MC38) mice/group pooled from 5 (D4M), 3 (D4M-S), or 3-4 (MC38) separate experiments. Bars depict the median values of the distribution. p values by Mann-Whitney U test. For **E**, the mean and SD of 8 (Iso) and 9 mice (αPD-1) in 2 separate experiments are shown. p value by Student’s *t*-test. **J-L.** Quantification of Treg (**J**), CD8 (**K**), and Th (**L**) cell numbers and percentage of CD45^+^ in lymph nodes draining D4M and D4M-S tumors. **M,N.** eTreg and CD44^hi^CD62L^−^ CD8 frequency (**M**) and Ki67 percentage (**N**) in tumor-draining lymph nodes. **O.** Foxp3, GITR, and ICOS MFI in eTregs from lymph nodes draining D4M or D4M-S melanomas. For **J-O**, n=25 (D4M) and 12-15 (D4M-S) mice/group pooled from 5 (D4M) or 3 (D4M-S) separate experiments. Bars depict the median values of the distribution. p values by Mann-Whitney U test. **P.** Experimental scheme to determine the role of PD-1 in cTreg, eTreg, and tumor Treg cells using bone marrow chimeras. **Q.** Treg frequency in the tumor-draining lymph node and MC38 tumors in radiation chimeras constructed to achieve 50% wt and 50% PD-1^-/-^ hematopoietic cells. n=3-5 mice/group. p values by Mann-Whitney U test. **R.** Treg quantification in bulk (Helmink, Gide) RNA sequencing datasets. p values by paired Student’s *t*-test.

**Figure S2.**
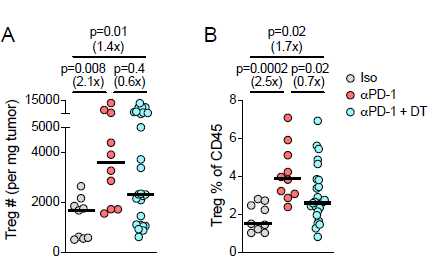
Treg abundance after partial DT ablation, related to Figure 2. Treg numbers per mg of tumor (**A**) and Treg percentage of CD45^+^ cells **(B)** in Foxp3^DTR^ mice bearing D4M-S tumors and treated as indicated. n=9 (Iso) 10 (αPD-1) and 27 (αPD-1 + DT) mice/group pooled from 2 separate experiments. Bars depict the median value of the distribution. p values by Mann-Whitney U test.

**Figure S3.**
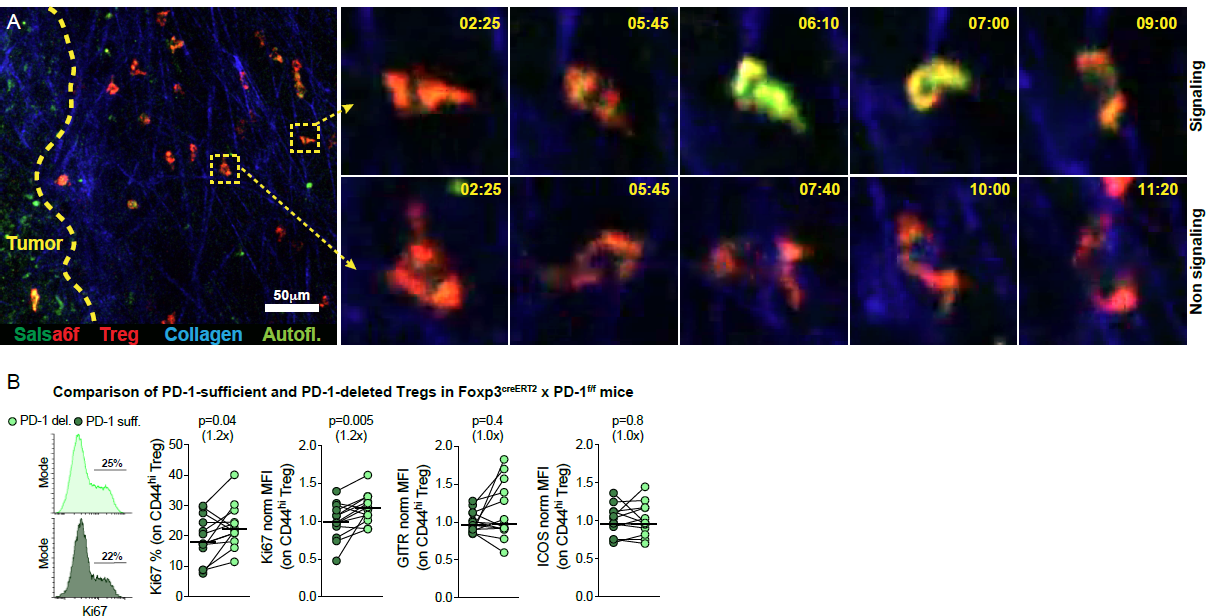
Treg imaging and effect of PD-1 deletion on Treg activation, related to Figure 3. **A.** Micrograph of the tumor-stroma border infiltrated by Salsa6f-expressing Tregs in Foxp3^creERT2^ x Rosa26^LSL-Salsa6f^ mice. The tumor is the weakly autofluorescent structure outlined on the left. Time sequences of representative non-signaling and signaling Tregs are shown on the right. Time in min: sec. **B.** Representative histograms of Ki67 and quantification of Ki67, GITR, and ICOS in PD-1-deleted or PD-1-sufficient tumor Tregs within tamoxifen-treated Foxp3^creERT2^ x PD-1^f/f^ mice (n=13, three separate experiments). Bars depict medians. p values by paired Wilcoxon test.

**Figure S4.**
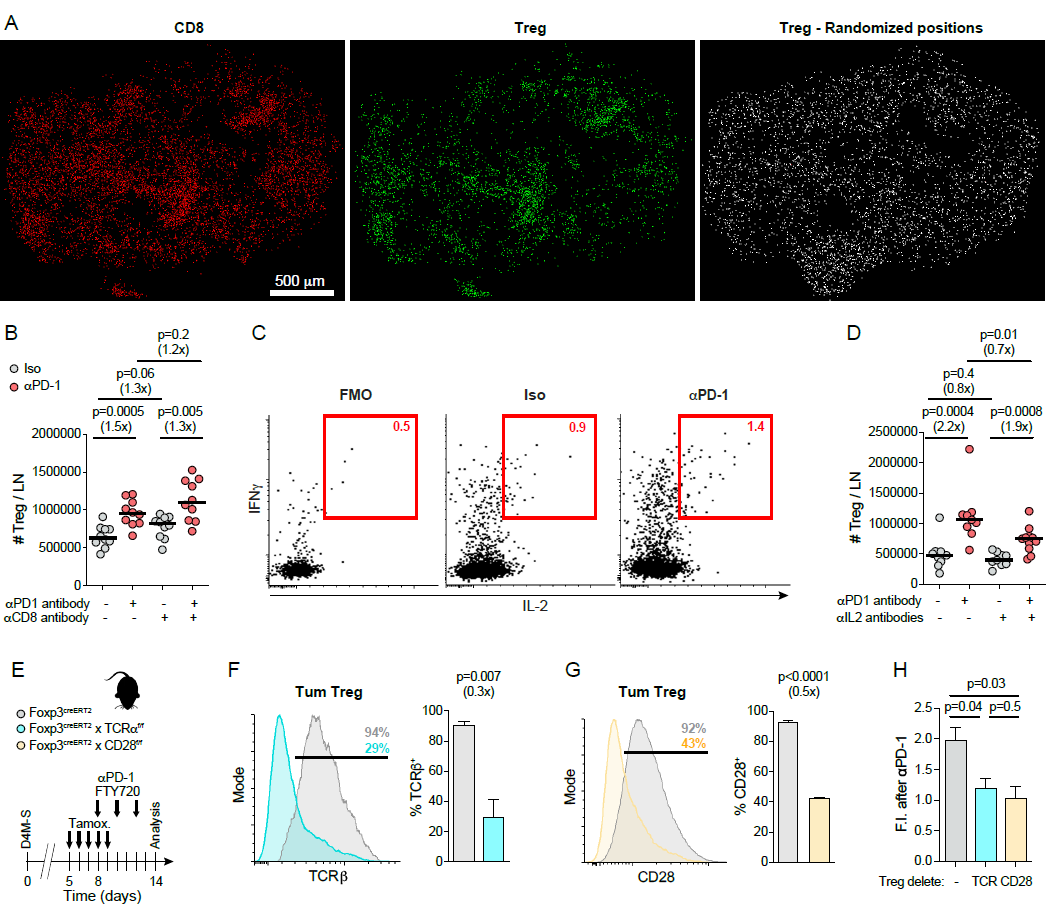
The role of CD8, TCR, CD28, and IL-2 in Treg accumulation after PD-1 blockade, related to Figure 4. **A.** Original positions of CD8 and Treg cells in the tumor shown in Figure 3A. Treg position after randomization is shown on the right. Each dot represents an individual cell. **B.** Number of Tregs per lymph node in mice bearing D4M-S tumors and treated or not with αPD-1 and CD8 depleting antibodies. **C.** Representative flow plots of IL-2 and IFNγ production in tumor CD8 cells after 8h restimulation with αCD3/αCD28 antibodies in the presence of brefeldin A. FMO = fluorescence-minus-one control. **D.** Number of Tregs per lymph node in mice bearing D4M-S tumors and treated or not with αPD-1 and IL-2 neutralizing antibodies. In **B-D**, n=9-11 mice per group in two independent experiments. Bars depict medians, and p value by Mann Whitney U test. **E.** Experimental scheme for specific TCR and CD28 deletion on Tregs in Foxp3^creERT2^ x TCRα^f/f^ or Foxp3^creERT2^ x CD28^f/f^ mice. **F, G.** Representative histogram and quantification of TCR (**F**) or CD28 (**G**) expression at sacrifice. **H.** Fold increase of tumor Treg percentages after PD-1 blockade. In Foxp3^creERT2^ x TCRα^f/f^ or Foxp3^creERT2^ x CD28^f/f^ mice, we evaluated the fold increase on Tregs negative for TCR or CD28, respectively. In **F-H**, mean ± SEM of 3-4 independent experiments is depicted. p values by Student’s *t*-test.

**Figure S5.**
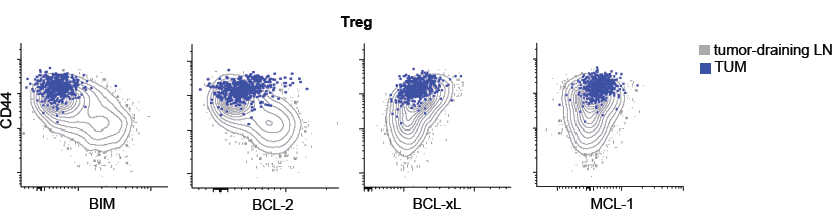
Expression of pro- and anti-apoptotic molecules in lymph nodes and tumors, related to Figure 5. Representative flow plots showing expression of the indicated molecules and CD44 in Tregs populating the draining lymph node (contours) or the D4M tumor (dots). One representative mouse out of 16 is depicted.

**Figure S6.**
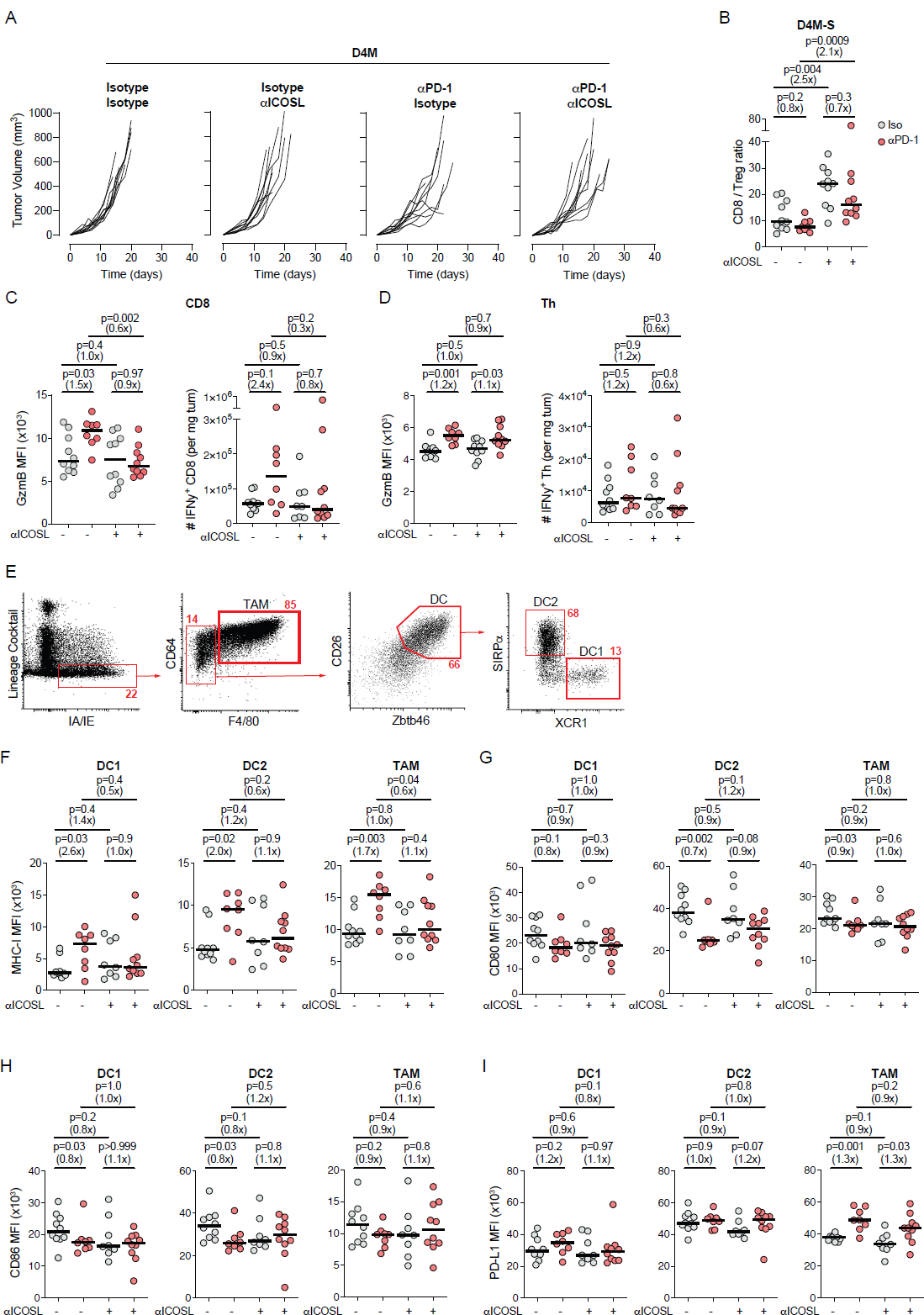
Effect of concomitant ICOSL / PD1 therapy on T cells and antigen-presenting cells, related to Figure 6. **A.** Growth curves of D4M tumors in C57BL/6 mice treated or not with concomitant αICOSL or αPD-1 antibodies. Each line represents one mouse. n=10 mice/group from 2 separate experiments. **B.** CD8/Treg ratio in D4M-S bearing mice treated with individual or combined ICOSL / PD-1 blockade. **C,D.** Granzyme B MFI and counts of IFNγ-producing cells in tumor CD8 (**C**) or Th (**D**). **E.** Gating strategy to identify DC1, DC2, and TAM cells in melanomas. **F-I.** MFI of MHC-I (**F**), CD80 (**G**), CD86 (**H**), and PD-L1 (**I**) in DC1, DC2, and TAM cells upon treatment with αICOSL and αPD-1 blockade alone or in combination. For **B-I**, n=8-10 mice/group pooled from 2 separate experiments. Bars depict medians. p values by Mann-Whitney U test.

## STAR METHODS

### KEY RESOURCES TABLE

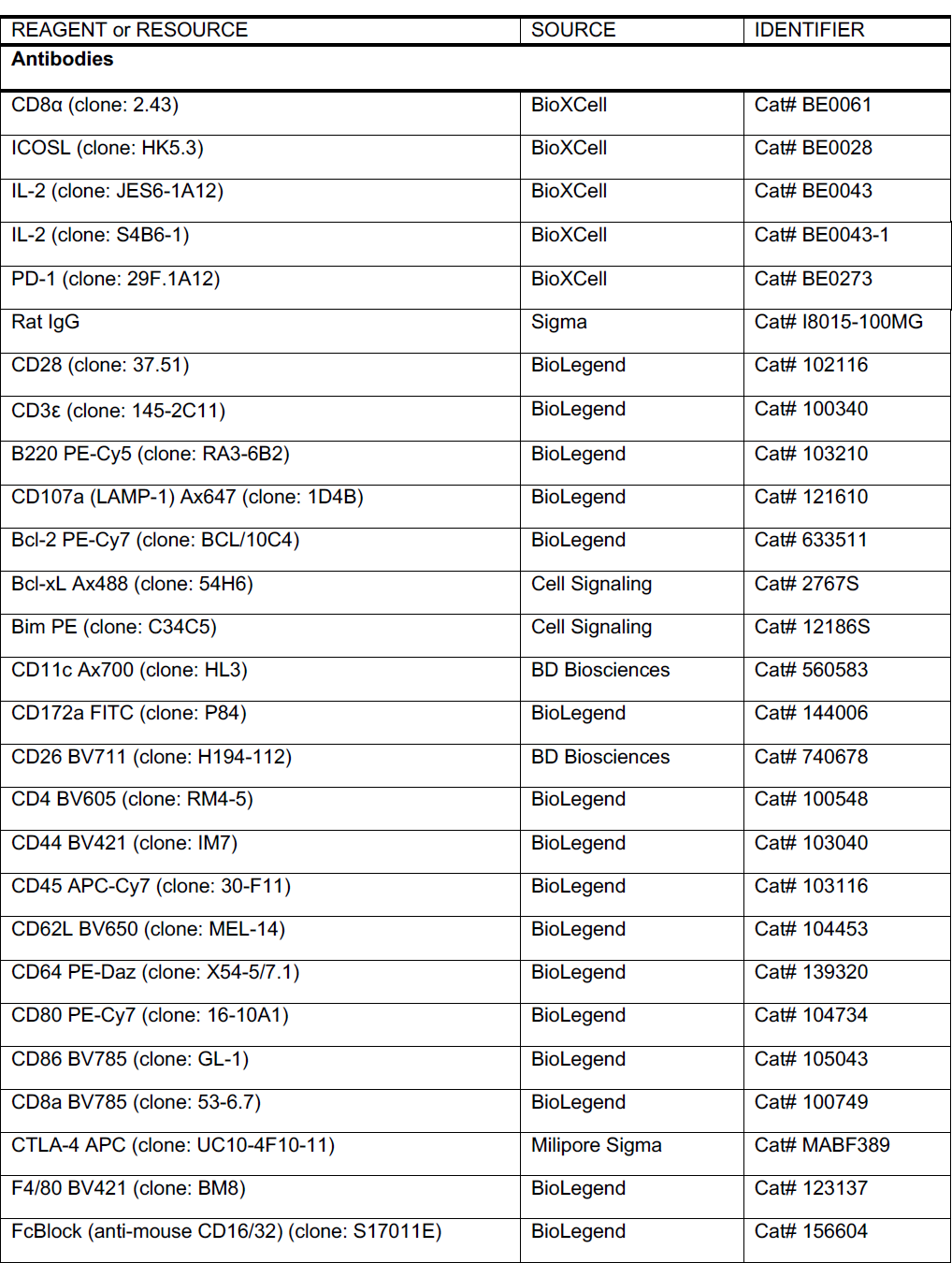
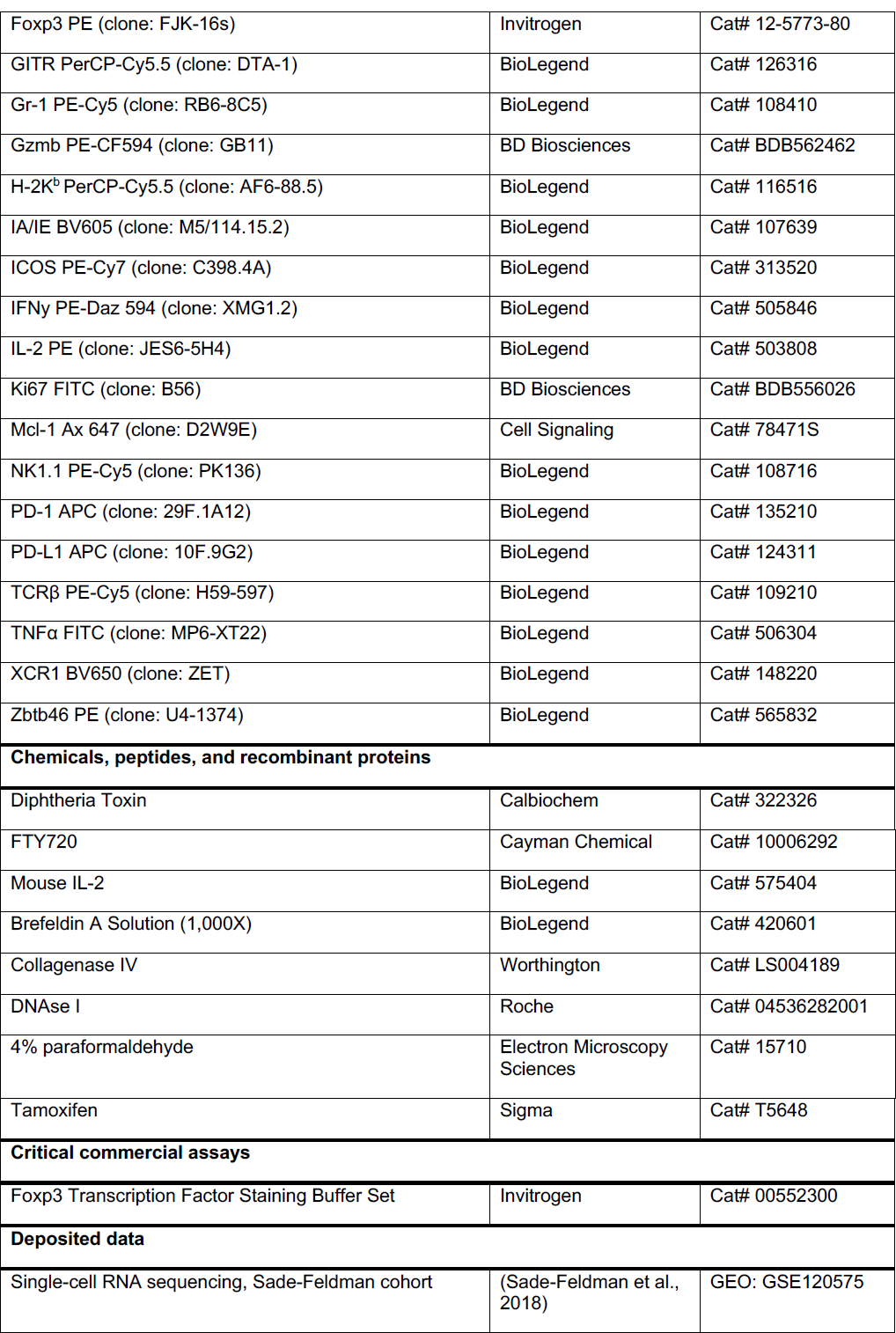
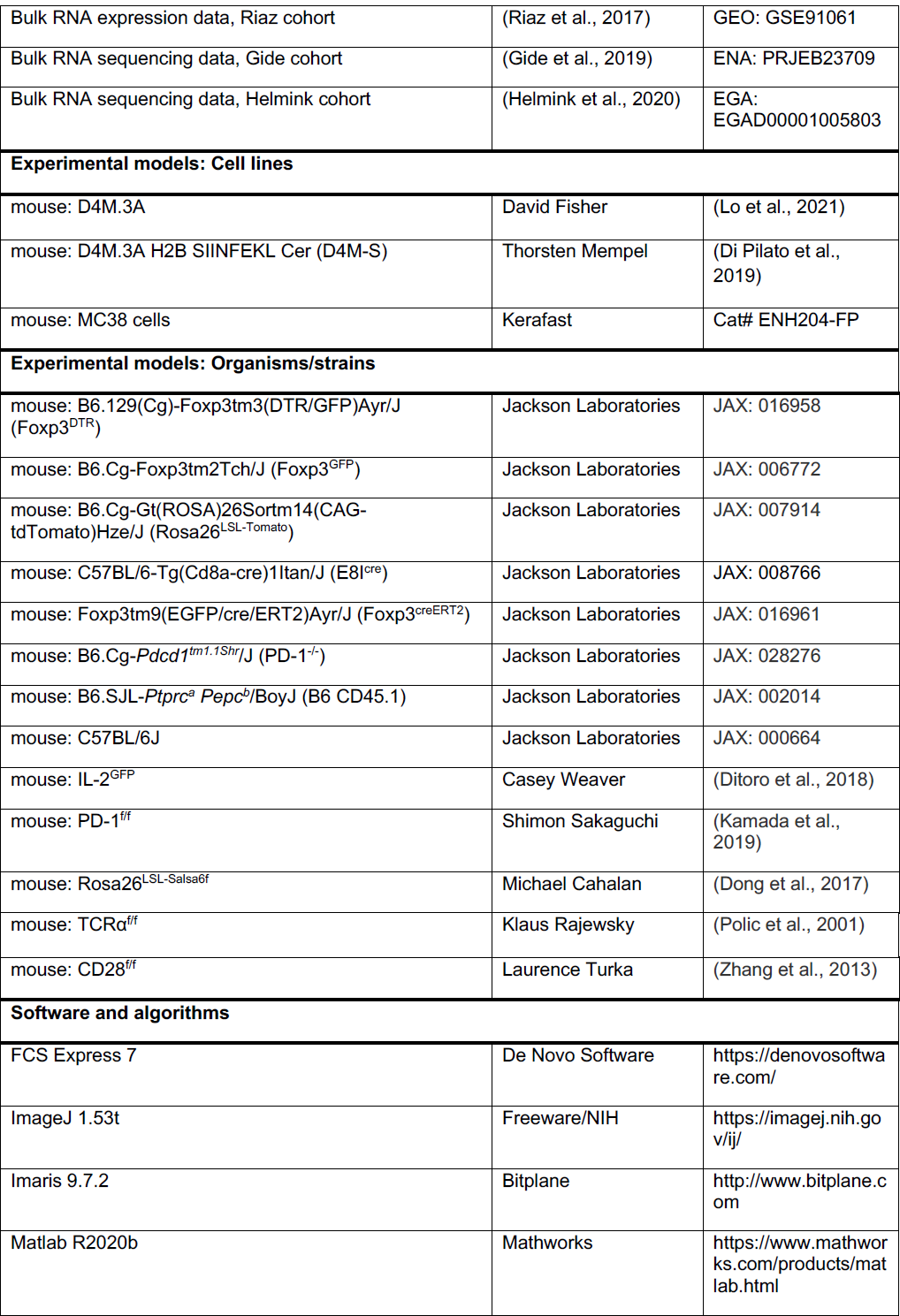
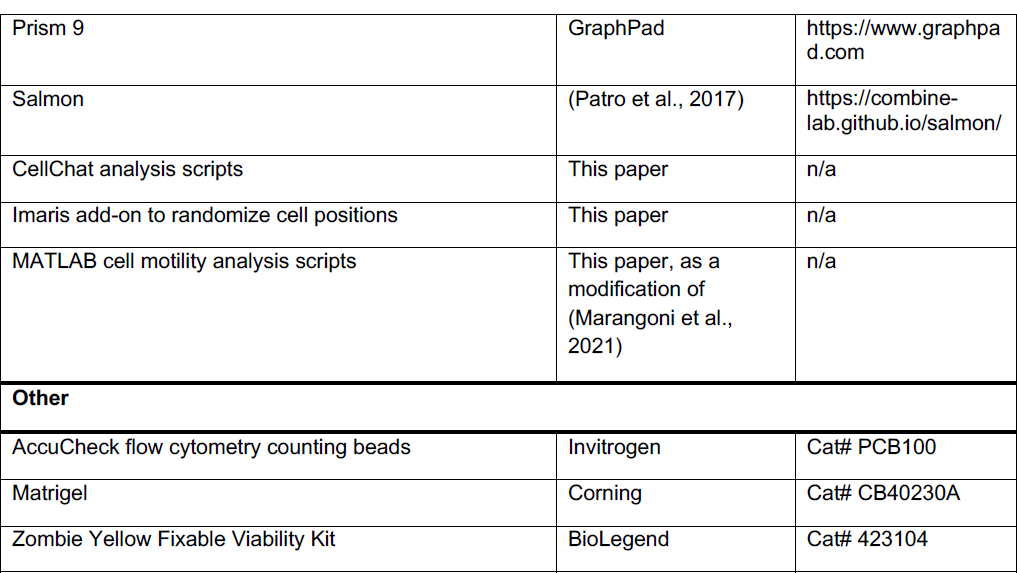

### RESOURCE AVAILABILITY

#### Lead contact

Further information and requests for resources and reagents should be directed to and will be fulfilled by the lead contact, Francesco Marangoni.

#### Materials availability

This study did not generate new unique reagents.

#### Data and code availability

- This paper analyzes existing, publicly available data. The accession numbers for the datasets are listed in the key resources table.
- Scripts for CellChat analysis, Imaris add-ons to randomize cell positions, and MATLAB Ca2+ signaling analysis scripts are available upon request.
- Any additional information required to reanalyze the data reported in this paper is available from the lead contact upon request.

## EXPERIMENTAL MODEL AND SUBJECT DETAILS

### Cells

D4M.3A (D4M) and D4M.3A H2B SIINFEKL Cer (D4M-S) melanoma cells were obtained from David Fisher and Thorsten Mempel, respectively, and grown in DMEM supplemented with 10% fetal calf serum (GeminiBio) under 37 °C / 5% CO_2_ conditions. These cell lines are derived from male mice (Jenkins et al., 2014) and have not been authenticated.

MC38 colon carcinoma cells were purchased from Kerafast (Cat# ENH204-FP) and grown in DMEM supplemented with 10% fetal calf serum (GeminiBio) under 37 °C / 5% CO_2_ conditions. MC38 cells are derived from female mice. This cell line has not been authenticated.

All cell lines were routinely tested for mycoplasma contamination and found negative.

### Mice

*E8i^cre^* (Maekawa et al., 2008), *Foxp3^creERT^*^2^ (Rubtsov et al., 2010), *Foxp3^DTR^* (Kim et al., 2007), *Foxp3^GFP^* (Haribhai et al., 2007), *Rosa26^LSL-Tomato^* (Madisen et al., 2010), PD-1^-/-^ (Keir et al., 2007), CD45.1 (Schluns et al., 2002), and C57BL/6 mice were purchased from The Jackson Laboratory. *CD28^f/f^*(Zhang et al., 2013), *IL-2^GFP^* (Ditoro et al., 2018), *PD-1^f/f^*(Kamada et al., 2019), *Rosa26^LSL-Salsa6f^* (Dong et al., 2017) and *TCRα^f/f^* (Polic et al., 2001) mice were obtained from the investigators who generated them. Mice were enrolled in experiments at 8-20 weeks of age. D4M cells derive from male mice and thus are artificially immunogenic in female recipients. To avoid this problem, we only implanted D4M and D4M-S melanomas in male mice. MC38 are of female origin and were studied in both male and female mice. In all cases, mice were bred, housed, enrolled in experiments, and euthanized according to protocols approved by the Institutional Animal Care and Use Committee (IACUC) of the University of California Irvine and the Massachusetts General Hospital.

## METHOD DETAILS

### Analysis of existing human datasets

Determination of Treg abundance in melanoma was carried out by selecting patients that i) were treated with PD-1 monotherapy only and ii) for whom pre-treatment and on-treatment data were available. Treg quantification in the Sade-Feldman dataset directly reflected the percentage for cluster G7 (Tregs) reported in Table S1 (Sade-Feldman et al., 2018). If a patient had multiple biopsies taken on-treatment, the Treg level was averaged. For the Huang dataset (Huang et al., 2019), we directly calculated the percentage of melanoma patients experiencing tumor Treg accumulation after PD-1 immunotherapy from Figure 4A of the original publication. To measure Treg abundance in bulk RNA sequencing datasets (Gide et al., 2019; Helmink et al., 2020; Riaz et al., 2017), we quantified the transcripts per million (TPM) for the GENCODE 32 GRCh38 genes using Salmon v1.9.0 (Patro et al., 2017). Treg cell abundance was inferred from the expression of FOXP3.

Cell-cell signaling pathways were analyzed using the R package CellChat (Jin et al., 2021). The CellChat package contains a manually curated database of ligand-receptor (L-R) interactions for the human and mouse species. CellChat receives as input the single-cell expression data as well as the cell type annotations for each cell and computes a “communication score” for every combination of sender cell type and receiver cell type, and for each L-R interaction. The count matrix and cell type annotations, split into pre-treatment and post-treatment groups, are taken from the original publication (Sade-Feldman et al., 2018). The original data were further filtered for treatment, and we maintained only patients that received PD-1 monotherapy (12 biopsies pre-treatment, 17 post-treatment, 10,609 cells total). The CellChat pipeline is first performed separately on the pre-treatment and post-treatment groups to compute the communication probabilities, as described in (Jin et al., 2021). CellChat also computes p-values for each interaction over each pair of cell types by performing permutation tests. Interactions were tested at a 5% significance level to identify cell types sending and receiving IL-2 signaling in the post-treatment group. Further analyses focused on the signaling from *TCF7*-expressing memory to Treg cells (group 10 to group 7 in (Sade-Feldman et al., 2018)), the only pair for which such signaling was identified. To quantify the change in IL-2 signaling in pre-treatment versus post-treatment conditions, we computed the fold change of the communication score and performed differential expression analysis on IL-2 in the *TCF7*-expressing memory T cells before and after therapy. We carried out the differential expression analysis using the Mann-Whitney U-test.

### Antibody treatment of tumor-bearing mice

Mice received a subcutaneous injection of 10^6^ D4M cells, 2×10^6^ D4M-S cells, or 10^6^ MC38 cells. D4M-S and MC38 cells were resuspended in Matrigel to facilitate engraftment. To maximize material for downstream analyses, we injected two tumors per mouse 1cm off the midline in both sides of the abdomen. In studies involving the deletion of floxed genes in Foxp3^creERT2^ models, tamoxifen treatment consisted of oral gavage (15 mg in 75 µl EtOH + 425 µl corn oil) on day 5 followed by four i.p. daily injections of 2 mg (in 10 µl EtOH + 40 µl corn oil). Six days before sacrifice, 200 µg αPD-1 (29F.1A12) was injected with 1mg/kg FTY720 i.p., and injections were repeated every other day. FTY720 blocks the egress of lymphocytes from secondary lymphoid organs, allowing us to study the effect of PD-1 blockade on an isolated tumor environment. IL-2 neutralization was performed by injecting 750µg of S4B6-1 and 750µg of JES6-1A12 i.v. every three days to block interaction with the α and β subunits of the IL2R. In studies using IL-2 immunocomplexes, 5 µg of IL-2 mAbs (JES6-1A12) were mixed with 0.5 µg per mouse of recombinant IL-2 and incubated at 37°C for 15 minutes. We injected IL-2 immunocomplexes every two days to trigger Treg expansion. αCD8 (2.43) and αICOSL (HK5.3) mAbs were administered every three days at a dose of 300 µg i.p. We injected rat IgG as an isotype control through the same route and at the same concentration as each antibody.

### Measurement of immunotherapy-treated tumors

In experiments to investigate the kinetics of tumor growth, we implanted only one tumor. For concomitant PD-1 and ICOSL blockade, 200 µg αPD-1 antibodies were administered every two days and 300 µg αICOSL antibodies every three days, beginning from day 13 after tumor implant. For sequential immunotherapy, αICOSL was injected on day 6 and 9 after tumor implantation, while αPD-1 mAbs were started on day 12 and given every two days. Tumors were measured three times a week with an electronic caliper, and tumor volume was estimated using the formula 0.5 x a x b^2^, where a is the maximum and b is the perpendicular tumor diameter. Immunotherapy administration was stopped when all mice either controlled the tumor or reached an endpoint as per our IACUC protocol. Conditions for sacrifice were: maximum diameter >15mm or both diameters >10mm.

### Diphtheria toxin treatment

In studies where Tregs were partially depleted, Foxp3^DTR^ mice received two D4M-S tumors and were treated with FTY720, αPD-1, and diphtheria toxin (Calbiochem) i.p. starting from day 12 and every other day after that. We titrated the amount of diphtheria toxin to 500pg/g to decrease Treg counts to the levels observed without PD-1 inhibition. Upon sacrifice, tumors were weighed and analyzed by flow cytometry.

### Bone marrow chimeras

To assess the role of PD-1 in the transition between cTreg to eTreg to tumor Treg cells, we generated radiation chimeras by injecting a mixture of CD45.2^+^ PD-1^-/-^ and CD45.1^+^ bone marrow into lethally irradiated (950 rad) CD45.1 mice. We titrated the PD-1^-/-^ and CD45.1^+^ bone marrow mix to produce a 1:1 ratio within lymph node cTreg cells. Mice were used in experiments two months after transplantation to ensure hematopoietic reconstitution.

### Flow cytometry

Tumor cell suspensions were prepared by digestion of finely minced tissue for 30 min at 37 °C using DMEM 10% FCS supplemented with 1.5 mg/ml Collagenase IV (Worthington) and 50 U/ml DNAse I (Roche). Tumor-draining lymph nodes were mechanically dissociated.

We stained 8×10^6^ cells except otherwise stated. Dead cells were stained through exposure to Zombie Yellow (1:200), diluted in PBS, for 15 min at 4 °C. We counted absolute cell numbers using AccuCheck flow cytometry counting beads (Invitrogen). Cells were subsequently treated with 5 µg/ml FcBlock for 10 min at 4 °C to decrease nonspecific Ab binding. Extracellular mAb staining was carried out at 4 °C for 20 minutes in FACS buffer (PBS 0.5% BSA, 2 mM EDTA). Cells were then permeabilized using the Foxp3 fixation-permeabilization buffer (Invitrogen), while intracellular mAb staining was performed at 4 °C for 30 min in Foxp3 wash buffer. We acquired the samples on a NovoCyte Quanteon flow cytometer and analyzed the data using FCS Express.

#### T cell activation panel

The panel to count T cells and analyze their activation included Zombie Yellow and αCD45, αCD8, αCD4, αFoxp3, αCD44, αCD62L, and mAbs against various activation markers including αKi67, αICOS, αGITR, αGzmb, αLAMP-1, and αCTLA-4.

#### Apoptosis regulators panel

We stained cells with a panel including Zombie Yellow and αCD45, αCD8, αCD4, αFoxp3, αCD44, αCD62L, αKi67, and mAbs against various controllers of apoptosis including αBim, αBcl-xL, αBcl-2, αMcl-1, and αICOS.

#### Cytokine panel

To measure cytokine production by T cells, 4 x 10^6^ live cells from tumor-draining lymph nodes or tumors were stimulated with plate-bound αCD3ε (10 µg/ml) and αCD28 (10 μg/ml) in the presence of Brefeldin A (5 μg/ml) for eight hours at 37 °C. Cells were finally stained using Zombie Yellow and αCD45, αCD8, αCD4, αFoxp3, αCD44, αIFNγ, αTNFα, and αIL-2 mAbs.

#### APC panel

To assess APC activation and numbers, we stained cells with a panel including Zombie Yellow, a lineage cocktail of mAbs against TCRβ, Gr-1, B220, and NK1.1, as well as αCD45, αCD64, αF4/80, αH-2K^b^ (MHC-I), αIA/IE (MHC-II), αCD26, αCD11c, αXCR1, αCD172a (SIRPα), αCD80, αCD86, αPD-L1, and αZbtb46 mAbs.

### Preparation of mice for F-IVM studies

We induced Salsa6f expression in Tregs by treating Foxp3^creERT2^ x Rosa26^LSL-Salsa6f^ mice with three 10 mg tamoxifen gavages (in 50 µl EtOH + 450 µl corn oil) spaced two days apart. Subsequently, mice were epilated by shaving and a brief application of hair remover cream. 7.5 x 10^5^ D4M-S cells (resuspended in 10µl of PBS) were injected in the center of the back, approximately 1 cm to the right of the midline. Seven to eight days after tumor injection, we surgically implanted a dorsal skinfold chamber (DSFC) such that the tumor was centered in the optical window of the DSFC. Analgesia was achieved by injecting 5 mg/kg carprofen s.c. pre-operatively and every 24 hours after that. We implemented two control groups that were later pooled due to similar results: mice imaged before administration of αPD-1 or 24h after injection of isotype control antibodies. These control groups were compared to mice imaged 24h after treatment with αPD-1.

### F-IVM time-lapse recordings

Mice were anesthetized with inhaled isoflurane. To prevent blurring artifacts due to respiratory and other physiologic movements, the DSFC was secured to the motorized stage using a custom-built platform. The DSFC was maintained at 37° ± 0.5°C utilizing a heating system (Warner Instruments) and a thermocouple-based temperature sensor placed next to the tissue. Mice were imaged using a Leica SP8 DIVE upright multiphoton microscope fitted with a Leica 25x water-immersion objective with a correction collar (HC IRAPO, NA = 1.0, WD = 2.6 mm). Insight X3 laser was tuned to 950 nm for optimal excitation of GCaMP6f and Tomato. For four-dimensional recordings of cell migration and signaling, stacks of 9 optical sections (X=350 μm, Y=350 μm; 512 x 512 pixels) with 4 μm z-spacing were acquired every 5 seconds to provide imaging volumes of 32 μm in depth per time point (voxel size 0.69 μm x 0.69 μm x 4 μm). Imaging depth was typically 30-120 μm below the DSFC glass. We detected emitted fluorescence and second harmonic as follows: PMT channel 1 bandwidth 465 – 486nm; HyD channel 2 bandwidth 490 – 545 nm; PMT channel 3 bandwidth 560 – 600nm. Datasets were imported in Imaris 9.7 (Bitplane) for analysis, generation of maximum intensity projections, and exporting as MPEG-4 movies.

### Analysis of cell motility and Salsa6f signaling

Image processing was performed using Imaris and Fiji plugins (version 1.53t). The threshold cutoff module was used to remove diffuse backgrounds for each channel, then a Gaussian smoothing of 0.8-pixel radius was applied to the entire image. Tomato (red channel) photobleaching was corrected using CorrectBleach plugin (Fiji) using the histogram matching method, and noise was reduced using the “Remove Outliers” filter with a radius of two pixels and two standard deviations. We tracked Tomato cells using the “spot” function of Imaris 9.7 (Bitplane) to obtain XYZ coordinates. Green channel intensities within these spots were used for Ca^2+^ measurements rather than typical Green/Red ratios (Salsa6f) to avoid potential red-channel intensity artifacts induced by the bleach correction algorithm. Ca^2+^ signaling was quantified through the mean fluorescence of the GFP (green, GCaMP6f) channel. We calculated the baseline green fluorescence for each track as a band centered on the 30^th^ percentile of fluorescence and having as extremes the difference between the 30^th^ percentile and the minimum fluorescence value. Thus, the upper limit of the baseline is (2 x 30^th^ percentile – minimum) of green channel fluorescence. This value was subtracted from all GFP fluorescence measurements to highlight fluorescence values above baseline. We then identified signaling track segments with the following characteristics: i) GFP signal above the baseline for at least 15 seconds; ii) segments shorter than one minute must have an AUC >1000; and iii) segments longer than one minute must have an AUC/duration ratio >800. These characteristics were established empirically so that automatically identified signaling segments matched with visually annotated ones on a subset of the data. We extracted the percentage of time a track is signaling, the maximum signaling peak fluorescence, signaling duration, and peak AUC using Matlab (Mathworks).

### Analysis of CD8:Treg colocalization in tumors

We implanted D4M-S tumors in Foxp3^GFP^ x E8I^cre^ x Rosa26^LSL-Tomato^ mice and harvested them after 11-14 days for tissue-wide imaging. Following euthanasia, tumors were carefully dissected and fixed onto a plastic coverslip using tissue adhesive (3M Vetbond), explants were fixed overnight in 4% paraformaldehyde, washed at least ten times with PBS, and imaged within 30 days. Individual 3D image stacks (X= 590 µm, Y= 590 µm, Z= 400-500 µm) were collected with a voxel size of 1.15 µm x 1.15 µm x 5 µm. Sequential excitation and detection were as follows. Insight X3 tuned to 950 nm, PMT channel 1 bandwidth 406 – 485nm. HyD channel 2 bandwidth 499 – 536nm, and PMT channel 3 bandwidth 560 – 620nm. 3D Image blocks were stitched using the Leica “merge” algorithm (10% overlap) to generate the montage images. To assess whether CD8 and Treg colocalized in montage images, we identified CD8 and Treg cells using the Imaris “spot” function and measured the distance of each Treg to the closest CD8 cell by the Imaris distance transformation algorithm. We then generated a 3D surface that includes all the tumor-associated T cells and randomized Treg positions within the 3D surface boundaries. The distance between randomized Treg cells and the closest CD8 was determined again.

## QUANTIFICATION AND STATISTICAL ANALYSIS

The numbers of individual cells, recordings, and animals analyzed are indicated in the figure legends. Two-tailed Mann-Whitney U test or Student’s *t*-test (in case of normal or lognormal distributions) were used to compare two groups. To analyze tumor growth curves, we used type II Anova followed by Holm’s post-test as indicated in (Enot et al., 2018). For categorical variables, we used the chi-squared test. All statistical tests were performed using Prism 9 (GraphPad). p values smaller than 0.05 were considered significant.

## SUPPLEMENTAL MOVIE GUIDE

**Supplemental Movie 1:**
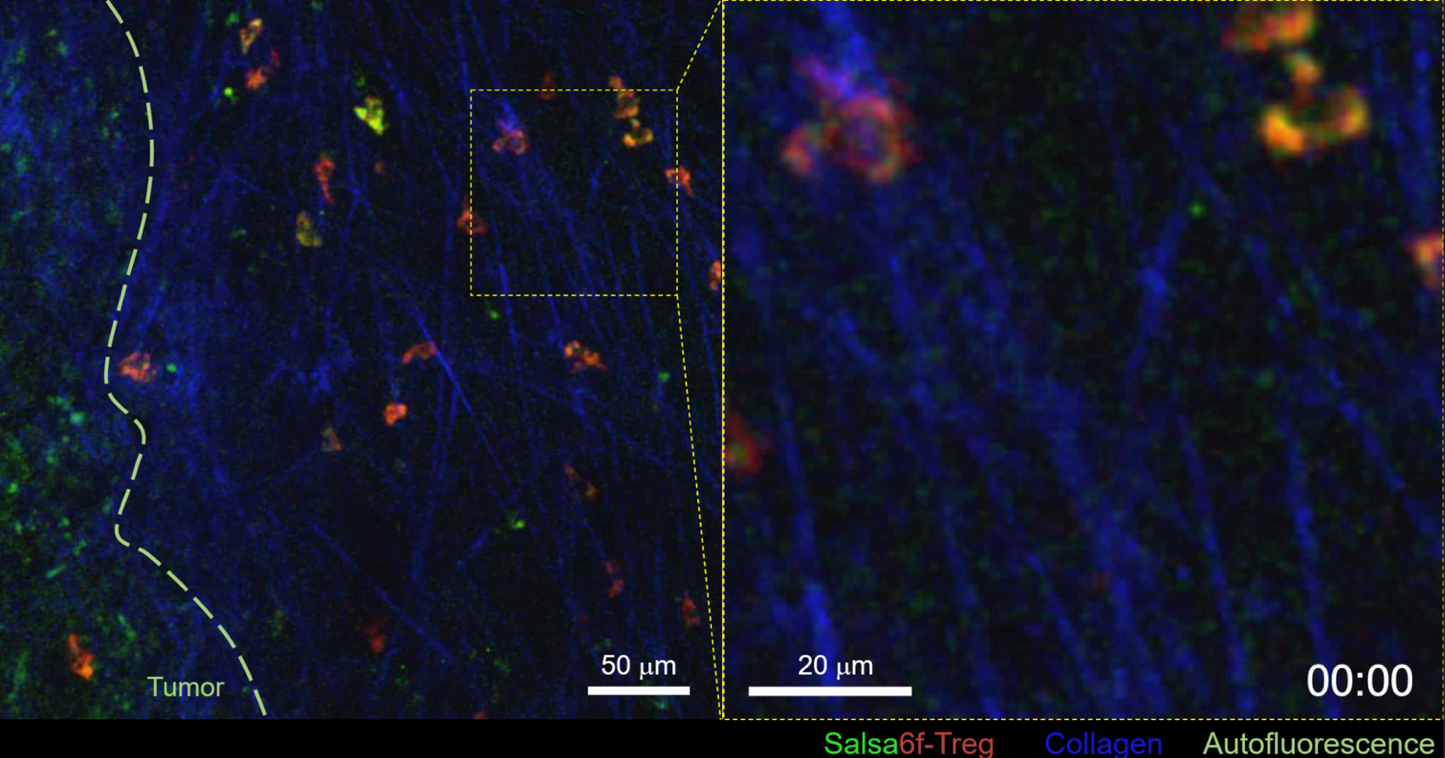
Visualization of Ca^2+^ transients in endogenous tumor Tregs (related to Figure 2) Left: Visualization of productive interactions of Salsa6f-expressing endogenous Treg cells. Tregs express Tomato (red fluorescence) constitutively, and flash green if they experience Ca^2+^ signaling. The dashed line depicts the tumor-stroma border. Right: Magnification of the region of interest highlighting Treg with Ca^2+^ dependent increase of green fluorescence. Time in min:sec.

**Supplemental Movie 2:**
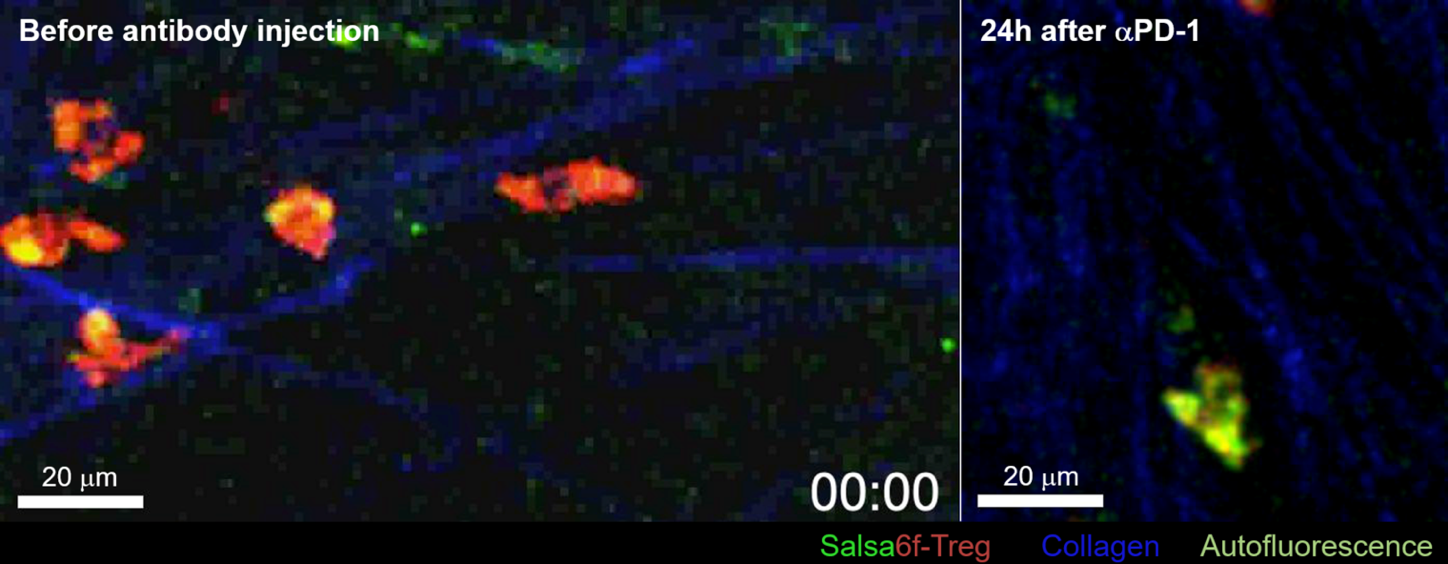
Ca^2+^ signaling in endogenous tumor Tregs before and after PD-1 blockade (related to Figure 2) Activation of Ca^2+^ signaling in endogenous tumor Treg cells before (left) or 24h after (right) administration of αPD-1 antibodies. Time in min:sec.

